# Skipper analysis of RNA-protein interactions highlights depletion of genetic variation in translation factor binding sites

**DOI:** 10.1101/2022.10.08.511447

**Authors:** Evan A. Boyle, Hsuan-Lin Her, Jasmine R. Mueller, Grady G. Nguyen, Gene W. Yeo

## Abstract

Technology for crosslinking and immunoprecipitation followed by sequencing (CLIP-seq) has identified the transcriptomic targets of hundreds of RNA-binding proteins in cells. To increase the power of existing and future CLIP-seq datasets, we introduce Skipper, an end-to-end workflow that converts unprocessed reads into annotated binding sites using an improved statistical framework. Compared to existing methods, Skipper on average calls 3.1-4.2 times more transcriptomic binding sites and sometimes >10 times more sites, providing deeper insight into post-transcriptional gene regulation. Skipper also calls binding to annotated repetitive elements and identifies bound elements for 99% of enhanced CLIP experiments. We perform nine translation factor enhanced CLIPs and apply Skipper to learn determinants of translation factor occupancy including transcript region, sequence, and subcellular localization. Furthermore, we observe depletion of genetic variation in occupied sites and nominate transcripts subject to selective constraint because of translation factor occupancy. Skipper offers fast, easy, customizable analysis of CLIP-seq data.

## Introduction

RNA-binding proteins (RBPs) conduct a vast array of essential functions in living cells. RNA synthesis, processing, modification, translation, and decay all require diverse RBPs that act in specific temporal, spatial, and cell type contexts^1,2^. Currently, crosslinking and immunoprecipitation followed by sequencing (CLIP-seq) methods are the gold standard for probing transcriptome-wide RNA-protein interactions in cells. Yet, CLIP-seq methods have continued to evolve and diversify^3,4^, requiring concomitant development of new tools for statistical modeling and data visualization.

CLIP-seq analysis must confront challenges inherent to analysis of both RNA-seq, where target sequences from transcript isoforms vary in expression by orders of magnitude, and ChIP-seq, where read signal aggregates into peaks against roughly even unbound chromatin background. CLIP-seq analysis tools generally identify binding sites by calling peaks^5–7^ or modeling positional enrichment^8–10^. Regardless of the approach taken, few tools attempt to test for binding to multi-mapping sequences and repetitive elements in the human transcriptome^7,11^ even though repetitive elements are in some cases the principal targets of RBPs^12^.

Peak calling approaches are ill prepared to handle diverse RBP binding modes. Results from peak calling approaches are susceptible to false negatives due to masking of intronic signal by exonic reads at exon-intron boundaries^13^, bias against calling peaks on low abundance transcripts^10^, and a mismatch between the length of bound regions and the bandwidth used for peak calling^9^. Apparent enrichment can vary either incrementally or abruptly over the course of a few nucleotides, but the sliding windows used to detect peaks are fixed in size. Even when binding profiles abide by expectations, reconciling partially overlapping peaks across samples and interpreting peaks that overlap multiple known transcripts is nontrivial.

Conversely, positional models can be underdetermined when binding sites are densely packed together and signals from distinct binding events overlap. More broadly, they require extensive parameterization that may be susceptible to biases related to CLIP-seq library fragment length or GC content. CLIP signal surrounding corresponding motifs often spans tens of nucleotides before decaying due to idiosyncratic patterns of RBP crosslinking to target sequences^5,8^. Cooperative and competitive RBP binding at nearby or overlapping binding sites can also alter RNA occupancy and regulatory outcomes^14–16^. Thus, interpretation of enrichment scores or binding affinity at nucleotide level resolution remains challenging.

Here, we introduce an end-to-end solution for analyzing CLIP-seq data (“Skipper”) that skips peak calling by tiling windows across annotated transcripts. Skipper processes both uniquely mapping and multi-mapping reads to report bound elements transcriptome-wide. Tiled windows and repetitive elements are tested for enrichment in immunoprecipitated over input samples using a beta-binomial distribution that accounts for overdispersion in read counts. We develop benchmarks for evaluating CLIP-seq data and compare Skipper to existing methods using eCLIP data available through the ENCODE Project website. Furthermore, we demonstrate the broad applicability of Skipper output by collecting new eCLIP data on a medley of translation factors and identifying selective constraint acting on translation factor occupancy as inferred from nucleotide sequence.

## Results

### Skipper allows rapid and adaptable analysis of eCLIP data

Before processing CLIP-seq read data, Skipper first tiles windows over annotated transcripts. Skipper optionally filters out genes that are not expressed in the cell type of interest to improve the accuracy of annotating overlapping features such as coding sequences, introns, or splice sites (Figure 1a). For any customizable set of transcript annotations, Skipper iterates over ranked features and transcript types to create variable length windows that do not traverse exon-intron junctions or gene boundaries. Skipper partitions the transcriptome into <100 nucleotide windows, corresponding to the length of library fragments generated by eCLIP (Figure 1a).

**Figure 1:**
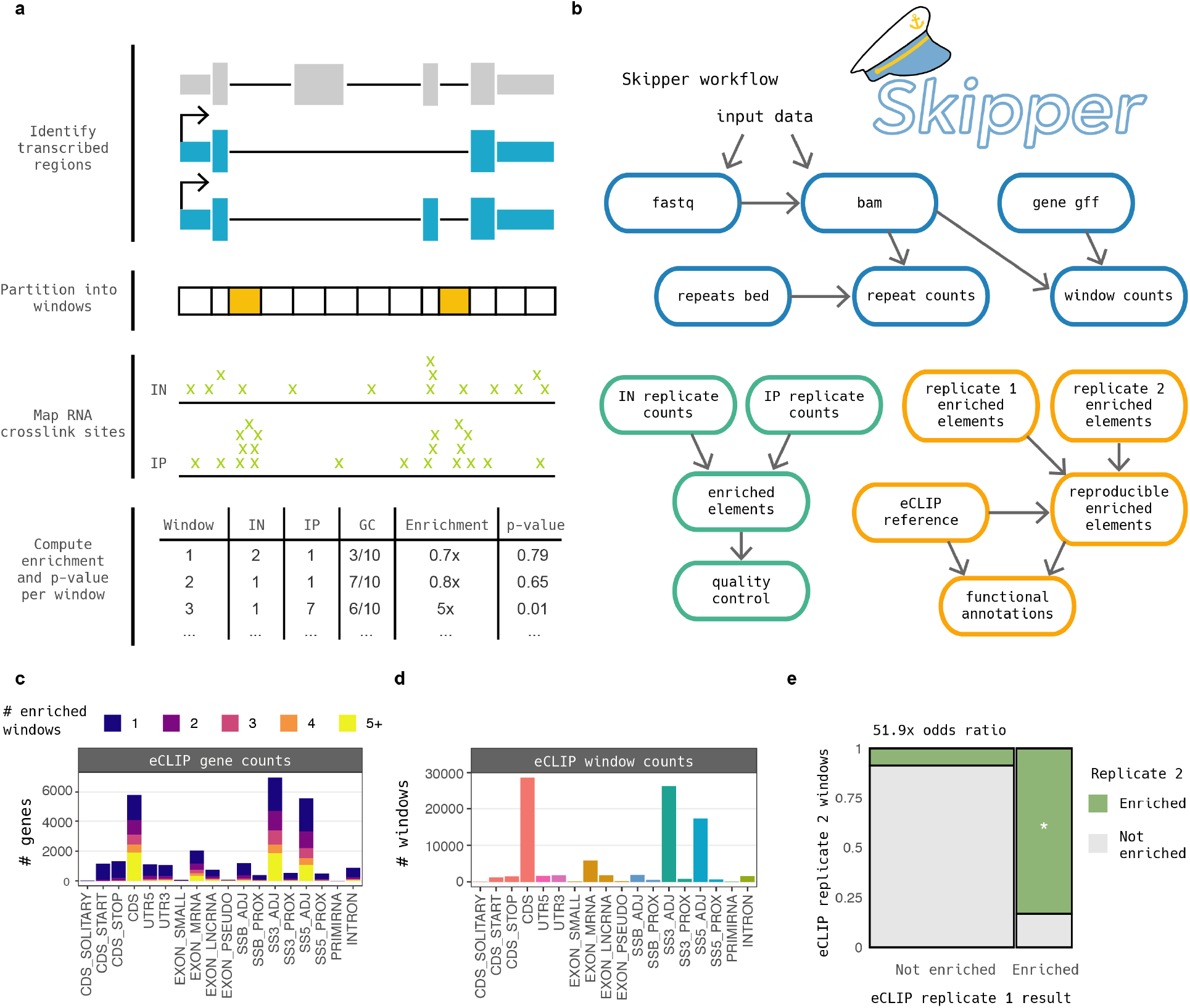
Calling RNA-protein interactions with Skipper. a) Illustration of Skipper binding calls starting from filtering transcript annotations (expressed transcripts in blue) to tiling the transcriptome with <100 nt windows, counting the number of crosslink sites (lime “x”s) per window for input (IN) and immunoprecipitated (IP) samples, stratifying by GC decile, and computing a p-value using the beta-binomial distribution (significant windows in orange). b) Outline of the full Skipper workflow. Each sample is processed into counts from one or more fastqs (blue). Enriched windows and repetitive elements are called (green). Reproducible elements are annotated and contextualized using publicly available eCLIP data (orange). c-e) Example output from running Skipper on AQR eCLIP in HepG2 including c) the number of bound genes, d) the number of enriched windows, and e) the concordance between replicates.

Skipper then begins sequence read processing (Figure 1b). Reads are trimmed and aligned, and multi-mapping reads are retained. Skipper quickly tallies the counts per window for each sample. Furthermore, reads are aggregated by repetitive element to consolidate multimapping sequences and permit quantification of binding to repetitive sequences. To test for enrichment in IP samples over input samples, a beta-binomial model is fit to the data. Overdispersion in read counts and GC bias are learned for transcriptomic windows and repetitive elements separately, and p-values are calculated from the fit beta-binomial distributions.

Extensive quality control summaries are generated for the resulting enriched elements. Output includes the number of bound genes (Figure 1c), number of enriched transcriptomic windows (Figure 1d), and the concordance between pairs of replicates (Figure 1e). After processing all replicates separately, reproducible enriched elements are ascertained for both transcriptomic windows and repetitive elements by selecting windows that pass a 20% false discovery in two or more replicates per experiment. Reproducible enriched elements undergo additional, optional, and customizable visualization and analysis.

Overall, Skipper’s total runtime is approximately 8-fold reduced compared to our previous peak calling pipeline based on CLIPper^17^ (Supplementary Figure 1a, Supplementary Table 1). We noted that overdispersion estimates and GC content vary across replicates of eCLIP experiments: adjusting for biases substantially changes the results of significance testing (Supplementary Figure 1b-c). In some cases, correcting for GC content radically alters the calling of enriched windows (Supplementary Figure 1d). Given that many RBPs favor AU-rich or GC-rich sequences, GC content differences are often confounded with true signal, but correcting at the level of approximately 100-nucleotide windows does not preclude recognition of short GC-rich motifs.

### Evaluating candidate binding sites across diverse RNA-binding proteins

To gain insight into Skipper output for diverse RBP binding profiles, we ran Skipper on all eCLIP fastqs available on the ENCODE project website.^17^ Across 219 eCLIP datasets, Skipper called an average of 21,310 reproducible enriched windows. We compared Skipper’s enriched window output to results from running Piranha^6^ on the same windows and overlapping significant CLIPper peaks with tested windows. For 72% of RNA-binding proteins, Skipper detected more enriched windows than both Piranha (average of 5,039 windows) and CLIPper (average of 6,904 windows). The disparity between Skipper and the other methods was greater for mRNA-binding RBPs than ncRNA-binding RBPs (Figure 2a, Supplementary Table 2).

**Figure 2:**
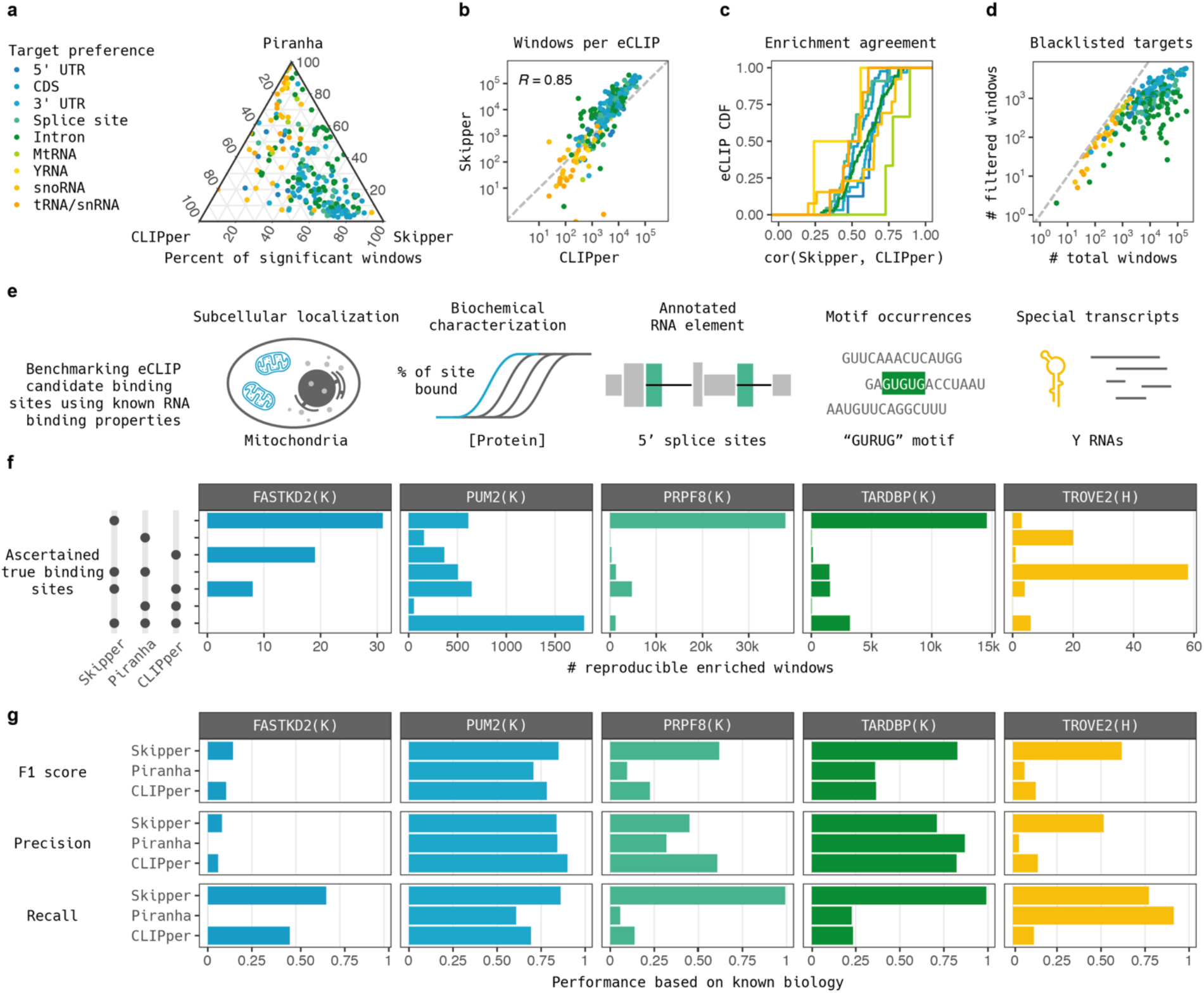
Benchmarking Skipper candidate binding sites. a) Ternary plot of the proportion of all enriched windows called by CLIPper, Piranha, or Skipper from ENCODE eCLIP data. b) ENCODE eCLIP experiments visualized by number of bound windows as called by CLIPper (x-axis) and Skipper (y-axis). c) Empirical CDF of the agreement in enrichment values between CLIPper and Skipper as measured by Pearson correlation, stratified by the target preference of the RNA-binding protein. d) Visualization of the effect of implementing a blacklist of nonspecifically bound sites. e) Ascertainment of true binding sites for example eCLIPs using mitochondrial subcellular localization for FASTKD2, biochemical characterization for PUM2, annotated 5′ splice sites for PRPF8, motif occurrence for TARDBP, and Y RNA identity for TROVE2. f) Counts of true binding sites per example eCLIP arranged in an upset plot. g) Per method summaries of precision, recall, and F1 score on Skipper performance in example eCLIPs.

Whether measured by number of enriched windows (Figure 2b) or agreement between observed enrichment values (Figure 2c), enriched windows detected by Skipper resembled CLIPper output. Skipper rarely increased the number of targets for RBPs that bind small noncoding RNAs like tRNAs, snoRNAs, snRNAs and Y RNAs: they are usually few in number but with robust immunoprecipitation enrichment.^5^ Effect sizes between the two methods were typically correlated around R = 0.6. RBPs that preferred mitochondrial transcripts, snoRNAs and 5′ UTRs were slightly more consistent (Pearson correlations of 0.77, 0.65, and 0.63, respectively) whereas splice sites, and tRNAs or snRNAs (Pearson correlations of 0.50 and 0.51) were more divergent (Figure 2c).

We noticed that some identified candidate binding sites were shared across many RBPs. To mitigate the potential for false positives due to biases in sequence alignment or library preparation, we derived a blacklist from the ENCODE project eCLIP data and filtered out the most common targets. RBPs that principally bind small noncoding RNAs (including SBDS, AARS, and SMNDC1) were most impacted by filtering: most Skipper enriched windows were removed and only a small number of snoRNA, tRNA, and snRNA windows remained (Figure 2d). By contrast, for intron and splice site binding proteins, only a small fraction of candidate binding sites were removed.

### Skipper vastly outperforms competing methods for calling RNA-protein interactions

Five distinct eCLIP experiments were selected for more rigorous evaluation of candidate binding sites: FASTKD2, which binds coding sequences; PUM2, which binds 3′ UTRs; PRPF8, which binds splice sites; TARDBP (or TDP-43), which binds introns; and TROVE2, which binds special regulatory RNAs known as Y RNAs. Annotations orthogonal to eCLIP data were further used to ascertain true binding sites, specifically mitochondrial transcript identity for FASTKD2, biochemically inferred binding affinity for PUM2^18^, annotated 5′ splice sites for PRPF8, presence of the GURUG motif for TARDBP, and Y RNA transcript identity for TROVE2 (Figure 2e).^19^

The number of ascertained true positive sites called by each method varied drastically across RBPs (Figure 2f, Supplementary Table 3). PUM2 was the most consistent across the three methods: nearly half of detected true binding sites were called by Skipper, CLIPper, and Piranha. For PRPF8, however, Skipper called seven times as many sites as CLIPper and ten times as many sites as Piranha. TARDBP exhibited a similar but attenuated trend, with Skipper calling more than three times more sites than CLIPper or Piranha. In the case of FASTKD2, mitochondrial transcripts were blacklisted and yielded zero binding sites under Piranha’s algorithm. TROVE2 was the only eCLIP dataset in which Skipper did not call the most putative binding sites.

Although Skipper reported the greatest number of ascertained true binding sites, the purity of true positives among called binding sites (the precision) is essential for the judging algorithm effectiveness. Somewhat surprisingly, the three methods exhibited similar precision despite large differences in the number of sites called (Figure 2g). Skipper attained the highest precision for FASTKD2 and TROVE2, Piranha for TARDBP, and CLIPper for PUM2 and PRPF8. Three cases exhibited particularly low precision: Piranha and CLIPper on TROVE2 and Piranha on FASTKD2 due to blacklisting and removal of mitochondrial transcripts.

Having established competitive levels of precision for Skipper, we further summarized Skipper performance. We calculated the percent of binding sites detected out of the union of ascertained true binding sites across all methods (the relative recall) and the harmonic mean of the precision and relative recall (the F1 score) for each eCLIP (Figure 2g). Skipper attained the highest F1 score across all examples. CLIPper output for FASTKD2 and PUM2 were the only cases where Skipper’s improvement did not exceed other methods by more than 10% as measured by F1 score. Because Skipper attained the greatest improvement in performance for TROVE2, we evaluated SLBP eCLIP, which binds a small class of specialized histone mRNA stem loops in the same way that TROVE2’s binds Y RNA motifs. Precision was comparable for all three methods, but Skipper again demonstrated the highest relative recall (Supplementary Figure 2). Examination of binding near alternative exons sensitive to knockdown revealed that Skipper also detected more RBFOX2 binding sites flanking knockdown-sensitive alternative exons than CLIPper (Supplementary Figure 3).

### A compendium of repetitive element binding

Our past work mapped reads to repetitive elements and examined information content across eCLIPs but did not report bound elements.^12^ With Skipper, we identify bound repetitive elements in eCLIP data using the same overdispersion and GC-content modeling framework. Nearly all (216/219) eCLIP datasets exhibited enrichment of repetitive elements (Figure 3a, Supplementary Table 4), whereas our past work limited its investigation of repetitive element enrichment to 65 eCLIP datasets.^12^ Bound repetitive elements achieved high specificity for known RNA templates, such as LARP7 and its binding partners 7SK and U6 (Figure 3b).

**Figure 3:**
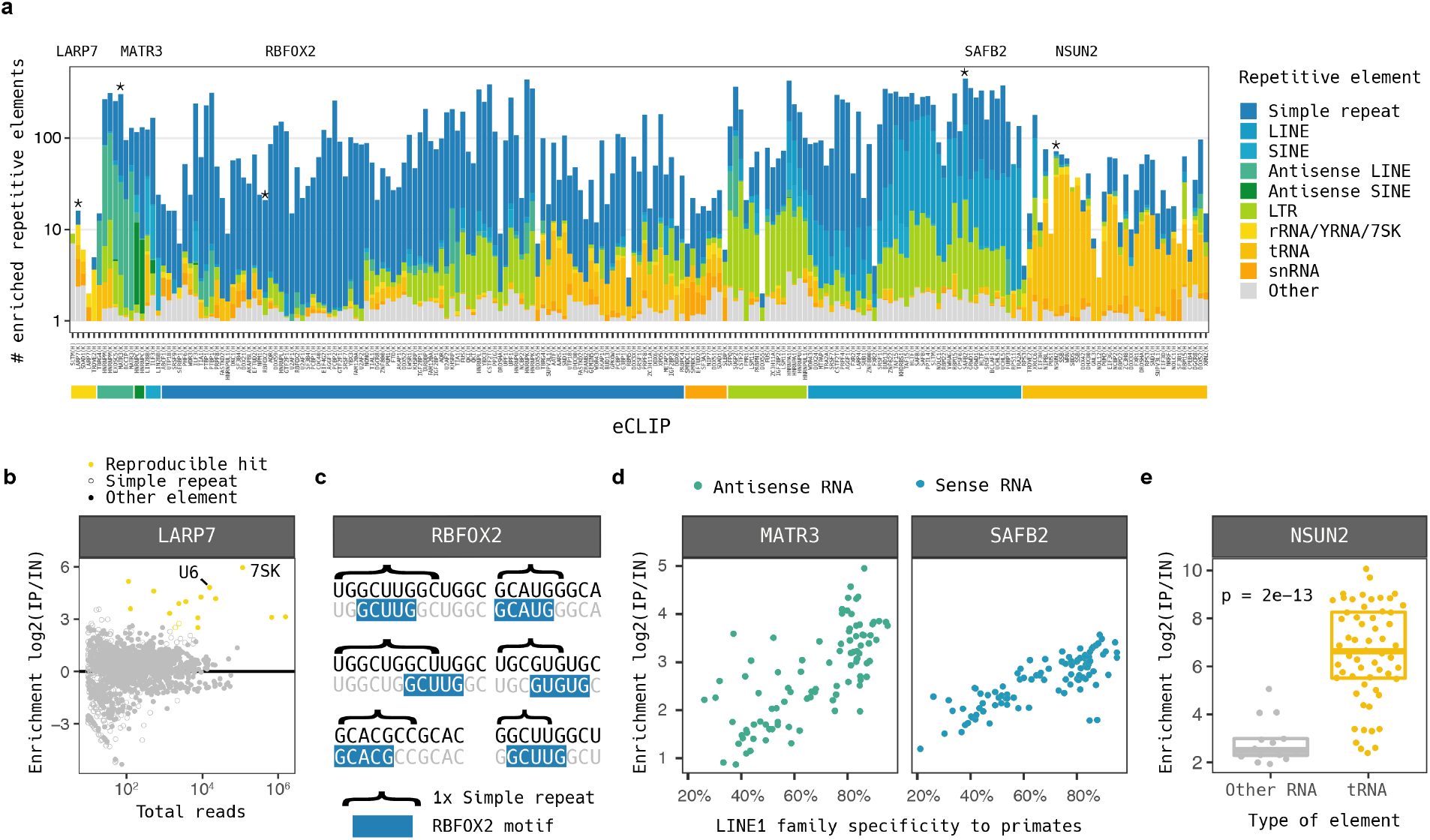
Skipper quantification of repetitive element binding. a) Counts of enriched repetitive elements per CLIP experiment, colored by type of repetitive element. Asterisks denote LARP7, RBFOX2, MATR3, SAFB2 and NSUN2, which are visualized in greater detail. b) LARP7 enrichments per element showing U6 and 7SK as most enriched among few other enriched sequences. c) Identification of RBFOX2 binding motifs contained within simple repeats. The top 6 enriched simple repeats are shown. d) Agreement between observed binding enrichment from Skipper and evolutionary age of LINE1 sequences in the human transcriptome for MATR3 (left) and SAFB2 (right), colored by strand of bound elements. e) Enrichment for tRNA-binding versus other identified elements for NSUN2.

Across all RBPs, the most frequently bound class of repetitive elements were simple repeats ranging from 1 to 12 nucleotides in length. Although we previously reported RBP motifs from reads transcriptome-wide, it has not been established whether long repeats of motifs show comparable specificity.^17^ We clustered mono-, di-, and tri-nucleotide repeat binding profiles and found both broad- and fine-scale patterns of selectivity (Supplementary Figure 4). GU-rich sequences were the most frequently bound repeats but exhibited diverse binding selectivity: some RBPs (e.g., LIN28B and LSM11) bound (UGG)n but not (GU)n repeats, whereas others (e.g., SRSF7 and TIAL1) bound (GU)n but not (UGG)n repeats. Even when RBPs bound both (UGG)n and (GU)n repeats, they differed in whether they bound (GUU)n repeats (e.g., HNRNPM and SUBP2), (AUG)n repeats (e.g. EFTUD2 and FAM120A), or neither (e.g., NCBP2 and FUS).

Some repeats exhibited a surprising level of specificity. Inspection of RBFOX2 simple repeats binding in K562 cells revealed that all 6 of the top simple repeats contained one of 8 established motifs directing canonical binding^20^ (Figure 3c), and out of all 16 simple repeats called, 11 contained one. QKI exhibited very high enrichment for (UAC)n repeats. QKI’s canonical binding motif UACUAACN_1-20_UAAY^21^ may commonly arise within (UAC)n repeats susceptible to mutation or transcription errors (Supplementary Figure 4). Finally, C homopolymers were bound by a small number of factors known to favor C-rich sequences including HNRNPK and PCBP2.

Other classes of bound repetitive elements generally agreed with known biology. MATR3, which extensively binds antisense LINE1 transcripts^22^, was among the top antisense LINE1 RBPs, and SAFB2, which represses LINE1 transcripts^23^, among sense LINE1 RBPs. In contrast to approaches that aggregate enrichment at the family level, Skipper tests binding to individual repetitive elements and robustly calculates enrichment even in cases of low read depth. We found that the reported enrichments strongly correlated with evolutionary age of the LINE1 element for both RBPs, and all bound LINE1 elements were of the expected strand (Figure 3d).^24,25^ Antisense SINE elements were bound most by HNRNPC^26^ and sense SINE elements by ILF3.^27^ We identified three main biological processes governing tRNA interactions: translation (EIF3G, RPS3, and SBDS), RNAi (SND1, DGCR8, and DROSHA), and tRNA modification (NSUN2, PUS1, and SSB). In some cases, these RBPs bound repetitive elements other than tRNAs with far lower enrichment than observed for tRNA targets (Figure 3e). Finally, snRNA-binding proteins included splicing regulators like SMNDC1 and EFTUD2.

A modest number of RBPs principally interacted with long terminal repeats (LTRs), which was not apparent from our previous enrichment filtering approach.^12^ Motif analysis with HOMER yielded mostly G- and GU-rich binding motifs. Notably, several of the RBPs in this class contain RGG motifs known to mediate RNA-protein interactions^28^: FUS, HNRNPA1, HNRNPU1, and HNRNPUL1. HNRNPK, which clustered with simple repeat-binding proteins, also contains an RGG motif and bound numerous LTR sequences. RGG domains are thought to mediate binding to G quadruplexes^29–31^, and recent work posits that stabilization of viral LTRs by RGG domain-containing proteins could play an important role in suppressing viral protein expression.^32,33^ Thus, in addition to their posited role in viral defense, RGG domain-containing proteins may also guard against reactivated endogenous retroviruses.

### Archetypes of alternative exon binding

Past studies have established enrichment of RBP binding sites at splice sites flanking alternative cassette exons and a depletion of RBP binding sites residing in alternative cassette exons.^17,34^ In aggregate, these binding profiles are appear to promote intermediate splicing; however, the identity of the bound RBP is critical in determining whether splice site usage is enhanced or silenced. Indeed, mutation of RBP binding sites induce splicing changes in accordance with the corresponding RBP’s distinct gene regulatory role.^35–38^

To examine alternative splicing regulation at the level of individual RBPs, we overlapped 5′ and 3′ splice sites flanking alternative cassette exons with the expanded set of enriched windows called by Skipper. We stratified the alternative exons by whether an RBP of interest bound the alternative splice site window, the constitutive splice site window, or both and searched for stereotyped RBP binding patterns (Figure 4a). Four patterns stood out, exemplified by SF3B4, RBM22, BUD13, and HNRNPC (Figure 4b). RBPs like SF3B4 principally bound 3′ splice site windows and favored constitutive 3′ splice site windows. RBPs like RBM22 principally bound 5′ splice site windows and favored both constitutive 5′ splice site windows and alternative 3′ splice site windows. RBPs like BUD13 commonly bound both 5′ and 3′ splice site windows with a constitutive bias for both splice site windows. Finally, RBPs like HNRNPC bound both 5′ and 3′ splice site windows with mild to moderate alternative window bias.

**Figure 4:**
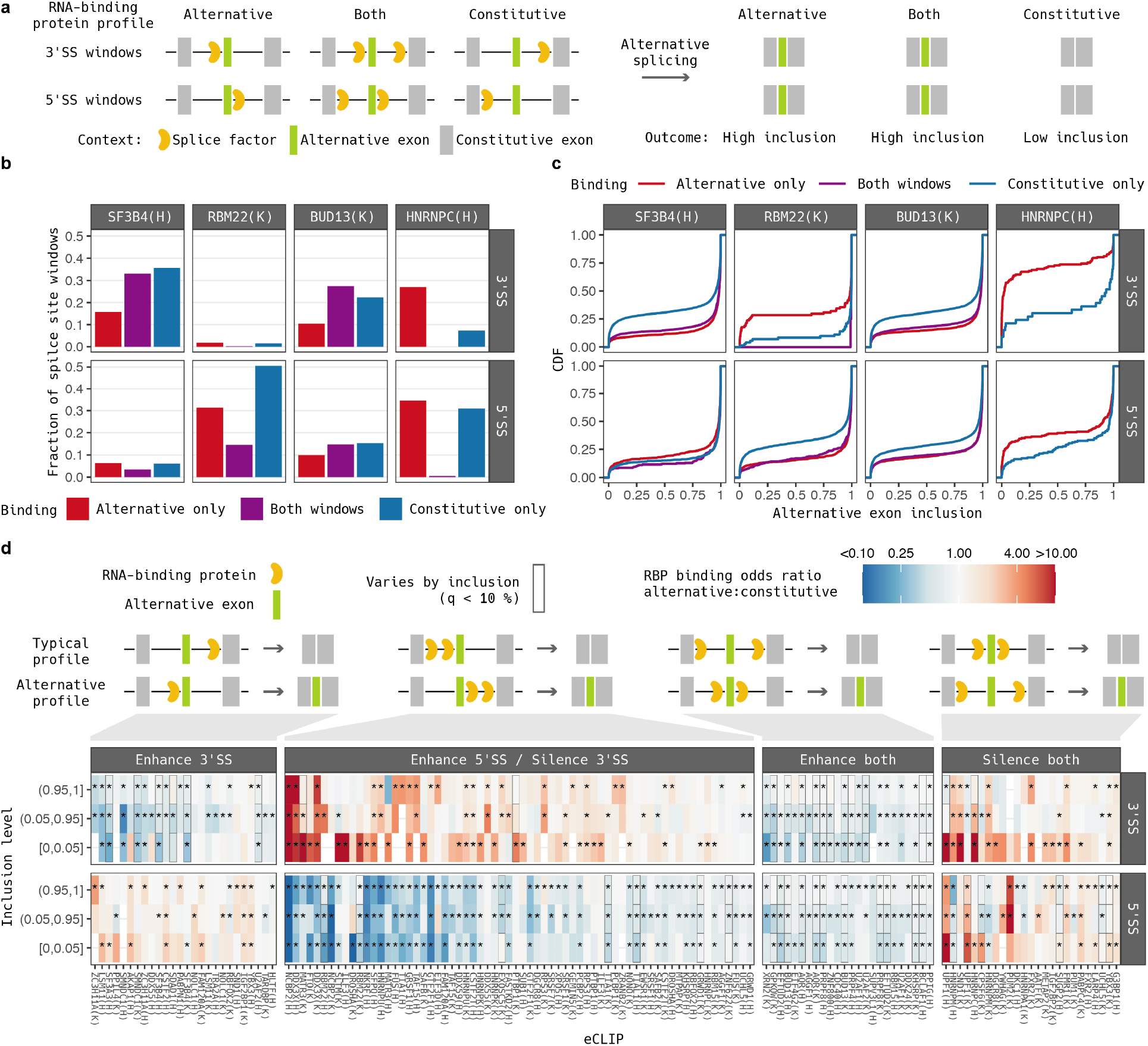
Archetypes of RNA-protein interactions near alternative cassette exons. a) Schematic of the approach taken for 3′ and 5′ splice site windows. RBP binding sites are tallied (left) at splice site windows flanking alternative (lime) and constitutive exons (gray) and associated to cassette exon inclusion levels (right). A general splice site-promoting factor is shown. b) Counts of alternative cassette exons where the indicated RNA-binding protein binds alternative (red), constitutive (blue) or both (purple) Skipper splice sites windows, for both 5′ and 3′ splice sites. c) Empirical cumulative distribution functions of alternative cassette exon inclusion stratified by the presence of binding sites in 5′ splice sites, 3′ splice sites, or both for the same four examples. d) Typical binding profiles of the 4 archetypes associated with skipping (top schematic) and alternative binding profiles associated with inclusion (bottom schematic). Heat shows alternative (red) and constitutive (blue) exon-biased RNA-binding proteins, significance in bias denoted by asterisks and significance in covariance with alternative exon inclusion denoted by boxed cells.

We then inspected the level of inclusion of skipped exons stratified by whether each RBP bound 5′ and/or 3′ splice site windows (Figure 4c). Alternative 3′ splice site binding by SF3B4 appeared to increase exon inclusion. Binding by RBM22 appeared to increase exon inclusion in alternative 5′ splice site windows and decrease exon inclusion in alternative 3′ splice site windows. In contrast, BUD13 binding appeared to increase cassette exon inclusion in the vicinity of either the 5′ or 3′ splice site. Conversely, HNRNPC binding appeared to decrease cassette exon inclusion when binding in the window containing either the 5′ or 3′ splice site.

Across all eCLIP experiments, most RBPs exhibited a binding profile that aligned with one of the four archetypes we described: 3′ splice site enhancing, 5′ and 3′ splice site enhancing, 5′ splice site enhancing / 3′ splice site silencing, and 5′ and 3′ splice site silencing (Figure 4d, Supplementary Table 5). Except for the 5′ splice site enhancing / 3′ splice site silencing archetype, greater exon inclusion levels usually corresponded to a shift from a typical binding profile linked to skipping to an alternative binding profile linked to inclusion. Among the 3′ splice site enhancing archetype members were well known 3′ splice site regulators (e.g. U2AF2 and SF3A3). The 5′ and 3′ splice site enhancing archetype included essential splicing factors like AQR and PRPF8. The 5′ and 3′ splice site silencing archetype included well established splice modulators like HNRNPM and SUGP2 as well as RBPs regulating mature mRNA stability, such as DGCR8, PUM1, CPSF6, and UPF1. In these cases, RBPs may be acting on overlapping alternative exons rather than intronic splicing elements.

Review of members of the 5′ splice site enhancing / 3′ splice site silencing archetype suggested that they were less likely to play a causal role in regulating alternative exon abundance (Figure 4d). Several RBP in this class did not canonically recognize splice junctions and instead bound 5′ UTRs (e.g., FTO, DDX3X, and NCBP2) or G- and GU-rich repeats (e.g., FUS, NKRF and GTF2F1). The binding profiles of these RBPs thus appeared to reflect a greater likelihood of residing within a 5′ UTR rather playing a regulatory role in RNA splicing or decay. Indeed, we also found instances of RBPs that favor 3′ constitutive splice sites and do not associate with exon inclusion (e.g., LSM11 and CSTF2), seeming to confirm that alternative splice site binding preferences do not always correspond to alternative RNA splicing or decay.

We repeated the binding site analysis using alternative 3′ splice sites and found that most significant differences in splice site occupancy reflected exon binding proteins that can bind alternative exons of mature mRNA in the cytosol, but not intronic windows in the nucleus (Supplementary figure 5, Supplementary Table 6). A small number of RBPs including PHF6, HNRNPL, and HNRNPK appeared to silence splicing in either alternative or constitutive windows. Conversely, essential splicing factors like AQR and PRPF8 promoted splicing in both alternative and constitutive windows. Finally, RBPs associated with regulation of alternative splicing, including RBFOX2, EFTUD2 and SMNDC1, bound the window flanking the alternative, longer exon regardless of inclusion level.

We crosschecked the 31 RBPs for which binding correlated with isoform abundance against RBPs known to regulate RNA splicing or decay. Members of our list of candidates were 3 times more likely to be known RNA splice factors than other RBPs that bound splice site windows (p = 0.01, Fisher’s exact test).^39–41^ Four other RBPs (YBX3, EXOSC5, UPF1, and MATR3) were annotated as playing a role in post-transcriptional gene regulation.

After removing the RBPs known to contribute to post-transcriptional regulation of gene expression, seven candidates for novel regulation of RNA splicing and decay remained: UCHL5, binding of which was recently linked to splicing changes via a multiplex splicing minigene screen^42^; GRWD1, binding of which is often disrupted by sQTLs^35^; and ZNF622, ZNF800, BCLAF1, DDX24, and SDAD1, for which binding has not been associated with RNA splicing or decay. ZNF622, ZNF800, BCLAF1, YBX3, and GRWD1 are putative transcriptional regulators that could influence RNA processing after transcription.^43^ DDX24 has been linked to RNA trafficking^44^, and SDAD1 is almost entirely unannotated.^45^ Notably, review of the repetitive element targets of UCHL5, GRWD1, ZNF622, ZNF800, BCLAF1, and DDX24 revealed primarily sense LINE1 transcripts (OR = 14 for candidate amidst all RBPs, p = 9*10^−5^, Fisher’s exact test, Figure 3a), suggesting that these factors may alter excision of introns containing LINEs.

### eCLIP of translation factors captures diverse molecular interactions

To demonstrate application of the Skipper pipeline, we generated new eCLIP data on a batch of translation factors with validated IP-grade antibodies: EIF2D, EIF2S2, EIF2B5, EIF3J, RPL35A, RPL29, RPS3A, RPS14, and RPS19 (Figure 5a). The binding preferences reported by Skipper varied drastically: RPL29 and EIF2B5 favored coding sequences, EIF3J and RPL35A favored 5′ UTRs, RPS14 and EIF2S2 favored snoRNAs, RPS3A favored tRNAs, EIF2D favored 3′ UTRs, and RPS19 favored introns (Figure 5b).

**Figure 5:**
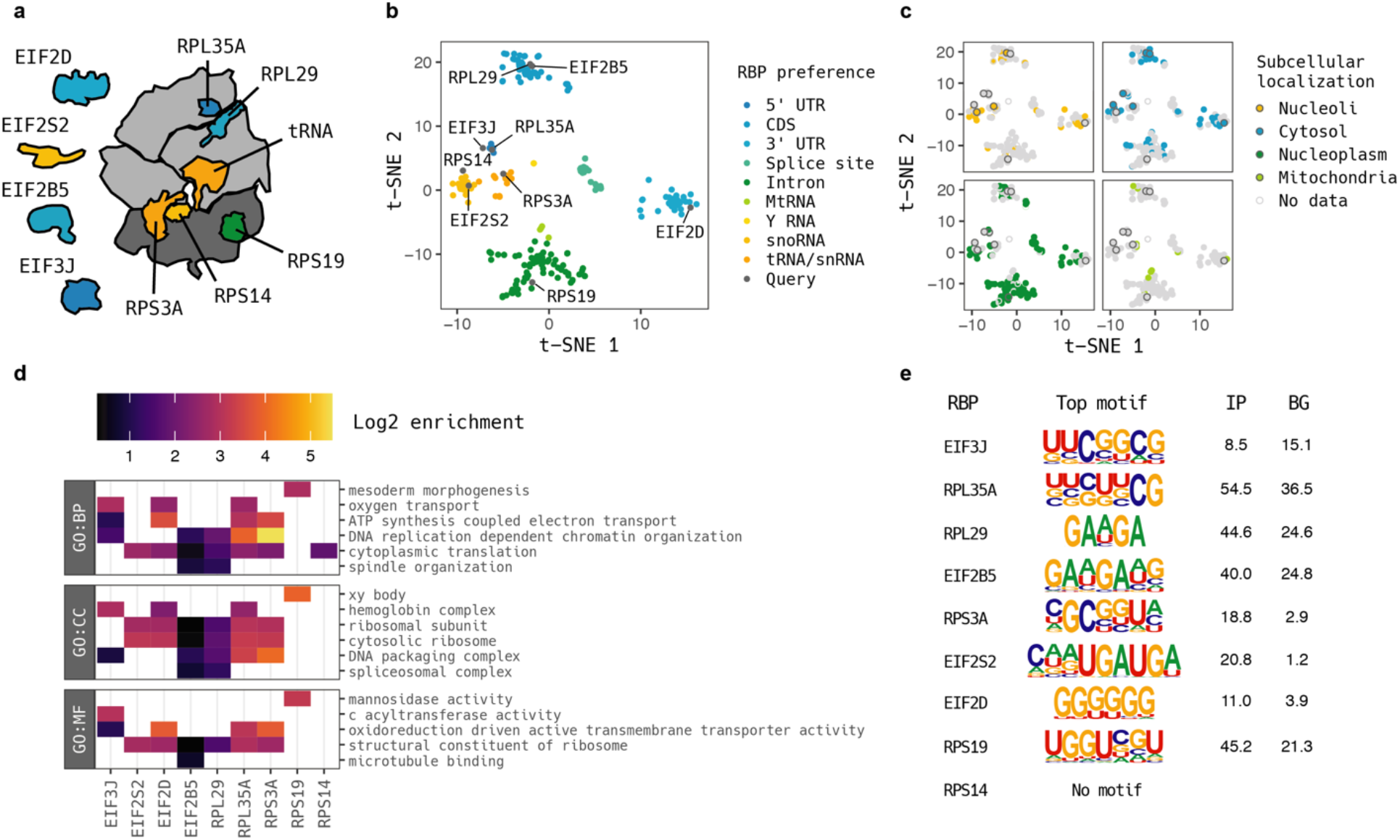
Determinants of translation factor occupancy. a) Illustration of profiled translation factors. b) Skipper t-SNE query of translation factors relative to clustered ENCODE eCLIP data colored by target preferences. c) Skipper t-SNE points recolored by subcellular localization to nucleoli (orange), cytosol (blue), nucleoplasm (dark green) and mitochondria (lime). d) Significant Skipper gene ontology enrichments for translation factors colored by log_2_ enrichment. At least the top cellular component, biological process, and molecular function term per eCLIP is shown. e) HOMER motif output for translation factor eCLIPs showing identified sequence motifs.

To interrogate the binding preferences of the selected translation factors further, we annotated the RBPs with their observed protein subcellular localization^46^ (Figure 5c). RPS19 localized overwhelmingly to the nucleoplasm, consistent with its strong intronic binding preference. EIF2S2 and RPS3A were detected in nucleoli, consistent with their preference for binding snoRNAs and tRNAs. The other translation factors localized to the cytosol, consistent with mRNA binding.

To discern which gene pathways were most enriched for translation factor binding, we performed a weighted gene ontology enrichment that tallied the number of enriched windows per term aggregating across the transcriptome (Figure 5d, Supplementary Table 7). Amongst the translation factors we profiled, the most commonly enriched term was “Cytoplasmic translation.” Related top terms include “Structural constituent of the ribosome”, “Cytosolic ribosome”, and “Ribosomal subunit”. Two other groups of genes were highly enriched: histone genes (“DNA packaging complex” and “DNA replication dependent chromatin organization”) and respiration (“ATP synthesis coupled electron transport” and “oxidoreduction driven active transmembrane transporter activity”).

We next ran HOMER to call sequence motifs underlying RBP binding. EIF2S2’s binding sites returned the snoRNA C box motif UGAUGA^47^, consistent with its snoRNA binding preference. RPL35A and EIF3J, both selective for 5′ UTR, exhibited motifs that reflect KYCKKCG binding. EIF2B5 and RPL29, both selective for coding sequences, both exhibited purine rich motifs. RPS19 yielded a GU-rich motif similar to other intron binding proteins: UGGUNGU. RPS3A was associated with GCGGU sequences, and EIF2D, known to regulate ribosome recycling near stop codons^48^, was associated with G-rich sequences in 3′ UTRs. RPS14 yielded no apparent motif.

Despite comprising many of the same protein complexes, different translation factors interacted with distinct sites in the human transcriptome. Binding profiles were largely distinct both from each other (Supplementary figure 6a) and from published ENCODE binding profiles (Supplementary figure 6b). Thus, translation factor binding profiles can reflect distinct biological processes that occur in varied subcellular compartments and exhibit distinct sequence preferences.

### Depletion of genetic variation nominates transcripts with constrained binding

Although our panel of translation factors exhibited distinct sequence preferences, this does not establish whether perturbation of these sequences would interfere with regulation of translation. Translation factor occupancy could conceivably be a passive readout of ribosome processivity with no functional significance. To test for constraint acting on sequence-driven translation factor occupancy, we trained a gapped kmer support vector machine (gkm-SVM) using LS-GKM on fixed 75-nt windows centered on signal overlapping enriched windows from Skipper and evaluated whether disruptive genetic variants were depleted in the gnoMAD genetic database.^34,35,49–51^.

gkm-SVMs for all translation factors but RPS14 exhibited significant separation between bound and control sites (Supplementary figure 7a). EIF2B5 and RPS19 target sites attained the greatest performance (area under the precision-recall curve, AUCPR, of 0.691 and 0.625) while the others exhibited low to moderate performance (AUCPR from 0.21 to 0.429). We moved forward with assessing potential constraint for all models but that of RPS14.

We queried gnoMAD for genetic variants in binding sites for each RBP and binned variants by allele frequency: singletons, very rare (<0.1%), rare (0.1-1%) or common (>1%) (Figure 6a). Reference and variant 75-nt sequences from Skipper binding sites were scored by the corresponding gkm-SVM to yield a delta score representing the predicted change in binding from the reference to the variant sequence. Delta scores were fit per transcript using linear regression against variant frequency bin as an ordinal variable. We interpreted greater slope in the linear regression as reflecting greater selective constraint on translation factor occupancy.

**Figure 6:**
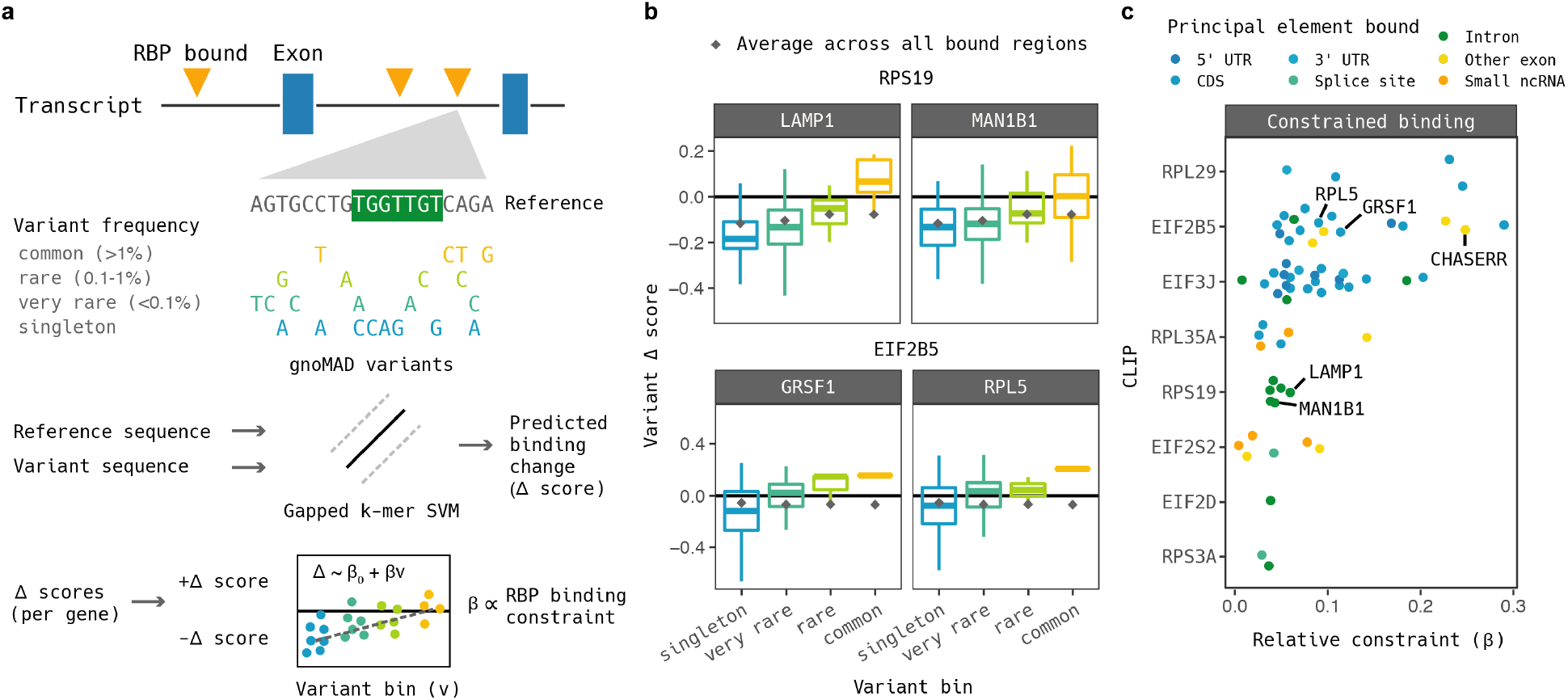
Identification of sequence-constrained translation factor occupied sites. a) Schematic of procedure for detecting constraint acting on occupied sites. gnoMAD variants are queried at all binding sites along a transcript. Genomic reference and variant sequences are scored using a gapped kmer support vector machine trained on Skipper binding calls. Variants are binned according to allele frequency and the difference between variant and reference predictions (the delta scores) are regressed against variant bin as an ordinal variable in a linear model. The slope is proportional to the selective constraint. b) Examples of transcripts with apparent sequence-constrained translation factor-occupied sites detected using RPS19 and EIF2B5 eCLIP. c) All 65 statistically significant constrained RNA-protein interactions colored by the primary feature type bound.

Some significant translation factor-transcript pairs were especially intriguing. Constrained RPS19 binding to *LAMP1* and *MAN1B1* transcripts was driven by multiple intronic windows. Singleton variants in binding sites had far lower delta scores than the transcriptome-wide average whereas variants above 0.1% frequency were far above (Figure 6b upper boxplots). The same trend was observed for EIF2B5 binding to coding sequences of *GRSF1* and *RPL5*, even though there was no transcriptome-wide trend disfavoring variants with lower delta scores (Figure 6b lower boxplots).

Overall, we detected 65 transcripts that were constrained for binding by a translation factor under a 10% false discovery rate (Figure 6c). Hits included long noncoding RNAs (e.g., *Chaserr*), demonstrating potential for discovery of noncanonical mechanisms of translation regulation. For RBPs that principally bound mRNAs, we also investigated characteristics that increased the likelihood of calling constrained transcript binding. RPL29, EIF2B5, EIF3J, and RPL35A were more likely to exhibit constrained binding in 5′ UTRs and less likely in introns and coding sequences (Supplementary Figure 7b). By contrast, transcripts containing RPS19 binding sites were much more likely to be constrained when binding sites occurred in introns than in coding regions. Thus, our nominated constrained binding events appear to recognize different gene regulatory roles acting on different transcript regions.

## Discussion

Our CLIP-seq processing tool Skipper offers fast, comprehensive analysis of CLIP-seq data with fully customizable reference and annotation files. Skipper matches or exceeds previous methods in precision and calls more than three times as many binding sites from eCLIP data available on the ENCODE project portal. With respect to interpretation of post-transcriptional functional consequences of genetic variation, Skipper increases the number of overlapping binding sites 2.5-fold for GTEx lead eQTLs and sQTLs (Supplementary Figure 8a-b), and 4-fold for lead 3′aQTLs (Supplementary Figure 8c-d). Unlike existing methods, Skipper also aggregates reads across instances of repetitive RNA elements and reports statistically enriched elements. Skipper’s automated visualizations expedite quality control of CLIP-seq data, and the corresponding tabular output provides a launching pad for exploration of RNA-protein interactions in high-throughput.

Evaluation of CLIP-seq data quality has overwhelmingly focused on detecting and quantifying motif enrichment^6–9,52^, but this approach overlooks many known determinants and consequences of RNA-protein interactions: subcellular localization that physically separates transcripts from RBPs, differences in binding affinity poorly reflected by position weight matrix motifs, essential RBP-mediated RNA processing including splicing, and specialized modifications or structural motifs that occur in specific classes of transcripts. We found that use of a size-matched input control improves performance dramatically in the case of the last group: in particular, Y RNAs by TROVE2 (Figure 2g), tRNAs by NSUN2 (Figure 3e), and histone mRNAs by SLBP (Supplementary Figure 2).^5^ The relatively small number of authentic targets for these RBPs increases vulnerability of analyses to false positives.

Similarly, the value of Skipper repetitive element output should not be overlooked. Binding to simple repeats was widespread (89.5% of RBPs exhibited 10-fold enrichment in binding to one or more simple repeats) but also distinct for each RBP (59.4% of RBPs possessed at least one simple repeat that no other RBP bound). Thus, our compendium should aid future efforts to use decoy RNAs^53^ to sequester RBPs at variable levels of specificity for diverse regulatory RBPs. Conversely, evaluation of potential decoy RBPs^54^ to block binding sites should be greatly facilitated using Skipper’s assignment of simple repeat preferences.

Investigation of RBP binding near alternative exons offers a complementary approach to post-transcriptional regulation. Characterizing RBP function using gene knockouts is often made impractical by the broad essentiality of RBP-regulated processes. While integration of eCLIP data with knockdown RNA-seq can define a causal link between binding and regulatory outcomes, siRNA knockdown is prone to off-target effects^55^, and imperfect calls for both RBP binding and isoform quantification (including differing GC bias and an insufficient number of informative reads) can limit the intersection across datasets^17^. Indeed, by correlating binding signal with alternative exon inclusion, we can infer many of the same relationships between RBP occupancy and RNA processing without collecting an independent knockdown RNA-seq dataset (Figure 4d). Our association of YBX3, ZNF622, ZNF800, BCLAF1, and GRWD1 differential binding to changes in isoform abundance suggests that the role of putative chromatin modulators in regulating RNA splicing and decay may be an underrecognized source of co- or post-transcriptional gene regulation.

Finally, our approach for detecting constraint acting on binding sites at the level of individual transcripts offers a roadmap for probing different layers of regulation by RNA-binding proteins. *Chaserr*, an essential lncRNA, serves as an example. Regulatory mechanisms for lncRNAs occur at myriad levels in gene regulatory networks from enhancer competition to post-translational modifications.^56^ *Chaserr* is thought regulate its neighboring gene CHD2 in cis, but the precise ways in which the CHASERR gene body fulfills its gene regulatory role are unknown.^57^ Selective constraint on sequences in loci occupied by translation factors suggests novel molecular mechanisms for regulation by *Chaserr* at the level of translation. The diverse types of transcripts, subcellular localizations, motifs, and RNA regions we identified under constraint by translation factors occupancy (Figure 5, Supplementary Figure 4b) reinforce the broad applicability of our approach.

Testing for transcript level constraint can also nominate mechanisms responsible for human disease. Mutations in RPS19 are the most common cause of Diamond-Blackfan anemia. The molecular pathophysiology of Diamond-Blackfan anemia entails defective rRNA maturation and impaired erythroid development, but why the erythroid lineage is predominantly affected over other cell types is unclear. Furthermore, patients exhibit considerable heterogeneity in clinical presentation that could point to co-occurring pathophysiology.^58^ Enrichment of disease variants occurring in RNA-binding protein binding sites is well documented, yet such work usually aggregates signal at the level of whole transcriptomes.^31,36,50^ By contrast, the transcripts with sequence-dependent RPS19 intron binding we identified under selective constraint could point to pathways that underlie erythroid susceptibility to mutations found in Diamond-Blackfan anemia patients as well as differing clinical presentations.

The transcript with the most constrained RPS19 binding, LAMP1, could conceivably play a role in Diamond-Black anemia pathophysiology. LAMP1 is one of the main constituents of lysosomal membranes. Erythroblasts rely on autophagy via lysosomes to eliminate organelles that impede erythrocyte maturation and function.^59^ One study found that knockout of factors essential for autophagy causes anemia in mice^60^, and an unbiased chemical screen revealed that induction of autophagy with the small molecule SMER28 enhanced erythropoiesis in induced pluripotent stem cells derived from Diamond-Blackfan anemia patients.^61^

As methodological and computational approaches to cataloging RNA-protein interactions continue to improve, understanding the functional significance of RBP binding will grow only more important. Determining whether an individual RBP binding site is under selective constraint remains challenging because a genetic variant’s predicted change in RBP binding depends on the precise nucleotide variant and its genomic context. Our results show that aggregating constraint at the level of transcripts is a well powered intermediate approach to generate hypotheses for functional transcript binding using publicly available datasets.

## Methods

### Tiling windows across the transcriptome

HepG2 and K562 total RNA-seq were downloaded from the encodeproject.org website. Transcript abundance was evaluated using Salmon^62^ per documented guidelines. GENCODE version 38 gene annotations were downloaded from the gencodegenes.org website, and transcripts with less than 1 transcript per million were filtered using the pyranges package in our custom script subset_gff.py for K562 and HepG2 cell lines separately.

We use the custom script parse_gff.R with a manual rankings of GENCODE accession types (roughly small noncoding RNAs first, then mRNAs, then lncRNAs, then pseudogenes) and feature types (ranges containing small noncoding RNA exons first, then both start and stop codons, start codons, stop codons, other CDS regions, 3′ UTRs, 5′ UTRs, noncoding isoforms of mRNA, lncRNA exons, 3′ and 5′ splice sites within 100 nucleotides, 3′ splice sites only, 5′ splice sites only, 3′ and 5′ splice sites within 500 nucleotides, 3′ splice sites only, 5′ splice sites only, primary transcript miRNAs, and finally introns) to tile windows. We iterate over ranked features and retrieve contiguous coordinates that are split into evenly sizes windows not exceeding 100 nucleotides in length. For example, a 50-nucleotide exon would yield a 50-nucleotide long window, but a 210-nucleotide exon would be split into three 70-nucleotide windows. The resulting disjoint windows are annotated with all overlapping transcripts and features and numbered uniquely.

### Skipper data processing

Skipper utilizes a single manifest of samples to preprocess, model, visualize, and summarize CLIP-seq data. For preprocessing, Skipper trims fastq files using skewer ^63^, cuts and pastes unimolecular identifiers using fastp^64^, aligns reads with STAR^65^, and deduplicates unimolecular identifiers using UMIcollapse^66^.

For beta-binomial modeling, counts of 5′ ends of reads per strand and GC content per tiled window and RepeatMasker element are calculated using SAMtools^67^ and bedtools^68^. Genomic positions that span multiple repetitive elements are ignored. Read counts are placed into ten GC contents bins for genome-mapped reads and twenty bins for repetitive elements where elements are binned in accordance with the average GC content of all instances of a particular repetitive element. A null overdispersion parameter (*rho*) and mean fold change parameter (*mu*) is estimated across bins by 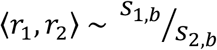 via the vglm function and beta-binomial family from the VGAM package ^69^, where r*i* is the counts per window or element in replicate *i*, and s*i,b* are the sums of counts in replicate *i* in GC bin *b*. With the overdispersion parameter estimated under the null, p-values are calculated for immunoprecipitated versus input samples with the *pbetabinom* function from VGAM. P-values less than 10^−12^ were replaced with 10^−12^ to address floating point imprecision. A false discovery rate is enforced by filtering windows for the sum of immunoprecipitated and input reads passing a dynamically determined threshold to maximize the number of hits and using the *p.adjust* function in R. Windows and repetitive elements that passed a 20% FDR in both replicates are called as reproducibly enriched, but reproducible enriched windows that were called in more than 17% of either HepG2 or K562 eCLIP samples are blacklisted and removed. Concordance between pairs of replicates is then assessed by Fisher’s Exact Test.

### Automated analysis of Skipper’s reproducible enriched windows

For clustering, transcriptomic windows counts are summarized as belonging to one of the following ranked categories: rRNA, snoRNA, snRNA, MtRNA, 7SK, Y RNA, and finally a combination of transcript type defined by the GENCODE gff and the feature type as defined above. Because of the large number of simple repeats, only repetitive elements enriched by at least 2.5 units of log_2_ fold change are included in clustering. Repeats that are not one of snRNAs, Y RNAs, tRNAs, LINE 1 sense or antisense, Alu sense or antisense, LTR, 7SK, or another LINE element are grouped together as “Other repetitive elements”. Each category is assigned an entropy contribution defined at 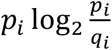 where p is the fraction of windows for category *i* in the queried CLIP sample and q is the fraction of windows in category *i* across all CLIPs. Reference data was clustered using Pearson correlation distance and the McQuitty agglomeration method via the *hclust* function in R. The nine classes of RBPs were created by cutting the tree into ten subgroups and reassigning a singleton clade. Each class was labeled according to the category with the greatest entropy contribution.

For motif calling, 75 nt control windows are created first by selecting a random window in the partitioned transcriptome of the same feature group – noncoding exons, mRNA exons, proximal introns (within 500 nt of exon-intron boundary), or distal introns – and then randomly selecting a center. Enriched windows are recentered on local maxima of binding enrichment (i.e. immunoprecipitated over input reads summed over 75 nt intervals) and scaled to a fixed 75 nt size. Multiple fixed 75 nt windows overlapping variable length enriched windows are iteratively selected until no local maxima remain or enrichment falls below the median across all positions. The findMotifsGenome.pl script from HOMER is run on the selected binding sites with the following options: -preparsedDir <directory> -size given -rna -nofacts -S 20 -len 5,6,7,8,9 -nlen 1 -bg <background coordinates>

For t-SNE visualization, CLIP-seq query data is pooled with ENCODE reference data and plotted in two dimensions using the Rtsne package.^70^

### Precision and relative recall calculations for CLIP-seq benchmarking

FASTKD2 enriched windows were considered true positives if they aligned to chrM and false positives otherwise ^17^. Scripts for assessing PUM2 binding affinity^18^ were modified to score arbitrary sequences. Enriched windows and matched control regions were extended 7 nt and scored. Scores above the 95^th^ percentile in the control regions were considered true positives and scores below the 50^th^ percentile true negatives. PRPF8 enriched windows were considered true positives if they were annotated as GENCODE 5′ splice site proximal (within 500 nt) and true negatives otherwise. TARDBP enriched windows were extended 5 nt and queried using bedtools nuc for the GURUG motif. Enriched windows containing the GURUG motif were considered true positives and true negatives otherwise^17^. TROVE2 enriched windows were considered true positives if they derived from Y RNAs and true negatives otherwise.^12^

IDR peaks are much narrower than the tiled transcriptomic windows used by Skipper and do not respect boundaries around UTRs and exon junctions. IDR peaks were reassigned the transcriptomic window that overlapped the center of the IDR peak. Thus, multiple IDR peaks within the same transcriptomic window were not double counted.

Because the full set of all true binding events is not known, the relative recall ignoring potential true positives outside the union of all ascertained true positives was used.

### RBFOX2 knockdown-sensitive exon analysis

RBFOX2 knockdown-sensitive exons were downloaded from Van Nostrand et al. 2016 by downloading source data to Supplementary Figure 10 (mislabeled as source data for Supplementary Figure 13 on the Nature Methods online article) and lifted over to GRCh38. Skipper windows or IDR peaks within 500 nucleotides of knockdown-sensitive exons were retrieved using bedtools flank and intersect commands and labeled with the corresponding exon SepScore.

### LINE1 evolutionary analysis

LINE1 specificity to primates was defined as the percent of individual instances of each LINE1 type that were novel to primates.^24^ The GC-corrected log_2_ enrichment per element reported by Skipper was then plotted against the specificity to primates.

### Evaluating RBP binding near alternative splice sites

Total RNA-seq for HepG2 and K562 cell lines was downloaded from the ENCODE Project website encodeproject.org. Alternative splicing was assessed using rMATS with the following options: --gtf gencode.v38.annotation.gtf --bi <STAR reference> -t paired --readLength 50 --od rmats -statoff. Alternative 5′ and 3′ splice sites were converted to BED format and intersected with all reproducible enriched windows called using ENCODE Project eCLIP data. Alternative exons with fewer than 20 total exon junction reads were discarded. Bias towards alternative or constitutive splice sites was calculated for each eCLIP experiment using the *binom.test* function with a probability of success (enriched windows overlapping the alternative splice site) of 0.5. Testing alternative versus constitutive splice site binding bias stratified by alternative exon usage was performed using the *chisq.test* function.

Known regulators of RNA splicing and decay were identified as belonging to the Gene Ontology terms “RNA splicing” and “Posttranscriptional regulation of gene expression”, plus CPSF6. TIAL1 and CPSF6 were not annotated with RNA splicing or post-transcriptional regulation of gene expression, but are well known to play those respective roles.^71,72^

### Assessing subcellular localization

RBP subcellular localization^46^ was downloaded from the Human Protein Atlas website proteinatlas.org. RBPs were noted for binding to the nucleoli, nucleoplasm, cytosol, or mitochondria in either the “Main location” or “Additional location” fields.

### eCLIP of translation factors

eCLIP was performed according to our published protocol^73^. eCLIPs were performed using antibodies from Bethyl Laboratories against EIF2B5 (A302-556A), EIF2D (A303-006A), EIF2S2 (A301-743A), EIF3J (A301-746A), RPS14 (A304-031A), RPL35A (A305-106A), RPS3A (A305-001A), RPL29 (A305-056A), and RPS19 (A304-002A) in K562 cells. The ribosome schematic was derived from entry 4V6X on PDB^74^.

### Selective constraint testing

LS-GKM was trained on 75-nt fixed windows centered on binding signal in translation factor eCLIPs^75^ and approximately 20,000 randomly sampled control windows (as for motif calling). Area under the precision recall curve was assessed via the ROCR R package^76^ using 5-fold cross validation estimates.

Genetic variants in gnoMAD overlapping 75-nt binding sites were queried remotely using bcftools looping across CLIP experiments and chromosomes: bcftools query -R $CLIP_bed -f ‘%CHROM\t%POS\t%ID\t%REF\t%ALT\t%INFO/AC\t%INFO/AN\n’ https://gnomad-public-us-east-1.s3.amazonaws.com/release/3.1.2/vcf/genomes/gnomad.genomes.v3.1.2.sites.${chromosome}.vcf.bgz

Variant fasta files were created from reference fastas by substituting single nucleotide variants one at a time. Delta scores were computed as the difference between gkm-SVM predictions on variant and reference fastas. Variants were placed into four bins: singletons, allele frequency < 0.1%, allele frequency between 0.1% and 1%, and allele frequency >1%, and delta scores were regressed against bin rank to determine relative constraint. To assess significance, delta scores were permuted within allele frequency bins 5000 times to create a null distribution of relative constraint – constrained transcripts were required to exceed the transcriptome-wide average trend. Instances in which no permutation exceeded the relative constraint of the observed values were replaced with 10^−4^, and the *p.adjust* function in R was used to enforce a 10% false discovery rate.

Enrichment for feature types in constrained transcripts was detected by Fisher’s Exact Test stratifying enriched window counts by whether the transcript passed statistical significance and whether the window derived from a particular feature type.

### Binding sites overlapping QTLs

eQTL and sQTL^37^ v8 datasets were downloaded from the GTEx portal. SNPs with the most significant p-value per gene or splice graph were retained as lead eQTL or sQTL, respectively. Lead 3′ alternate polyadenylation QTL (3aQTL)^77^ were downloaded from Synapse ID syn22131046. Polyadenylation sites^78^ were downloaded from GEO ID GSE138197. Skipper enriched windows and windows containing IDR peaks were intersected SNP positions using bedtools.

## Data availability

ENCODE eCLIP fastqs and CLIPper processed data are available on the ENCODE Project website: https://www.encodeproject.org/encore-matrix/?type=Experiment&status=released&internal_tags=ENCORE

Translation eCLIP raw fastqs and reproducible enriched window output are available at GEO: GSE213867.

Additional summary data for Skipper output are available on Figshare: 10.6084/m9.figshare.21206009

## Code availability

The Skipper pipeline including example input are available under a BSD license at https://github.com/YeoLab/skipper/ and deposited on Figshare at 10.6084/m9.figshare.21272991

## Acknowledgements

This work was supported by grants R01 HG004659, R01 HG011864, RF1 MH126719 and U41/U24 HG009889 to G.W.Y. G.W.Y. is supported by an Allen Distinguished Investigator Award, a Paul G. Allen Frontiers Group advised grant of the Paul G. Allen Family Foundation. G.W.Y. is a cofounder, member of the board of directors, equity holder, and paid consultant for Locanabio and Eclipse BioInnovations and a distinguished visiting professor at the National University of Singapore. The terms of these arrangements have been reviewed and approved by the University of California San Diego in accordance with its conflict-of-interest policies. E.A.B. is a Helen Hay Whitney Foundation Fellow. Data processing was performed on the Triton Shared Computing Cluster. This publication includes data generated at the UC San Diego IGM Genomics Center utilizing an Illumina NovaSeq 6000 that was purchased with funding from a National Institutes of Health SIG grant (#S10 OD026929). Data was also generated by the Sequencing Core Facility at the La Jolla Institute funded by National Institutes of Health SIG grant #S10 OD025052. The authors declare no other competing interests.

eQTL and sQTL data were generated by the Genotype-Tissue Expression (GTEx) Project and retrieved from the GTEx Portal in January 2021. Diego Calderon, Ryan Marina, and Katherine Rothamel provided helpful comments on the Skipper pipeline, text, and figures. Steven Blue and Brian Yee assisted with eCLIP metadata curation.

## Author contributions

E.A.B. conceived of the study, implemented the Skipper pipeline, analyzed data, visualized results, and wrote the manuscript. H.H. compiled a draft of Skipper preprocessing steps and reviewed eCLIP data quality measures. J.R.M. generated ribosome protein eCLIP data. G.G.N. generated EIF eCLIP data. G.W.Y. supervised the study and edited the manuscript.

---

**Supplementary Figure 1:**
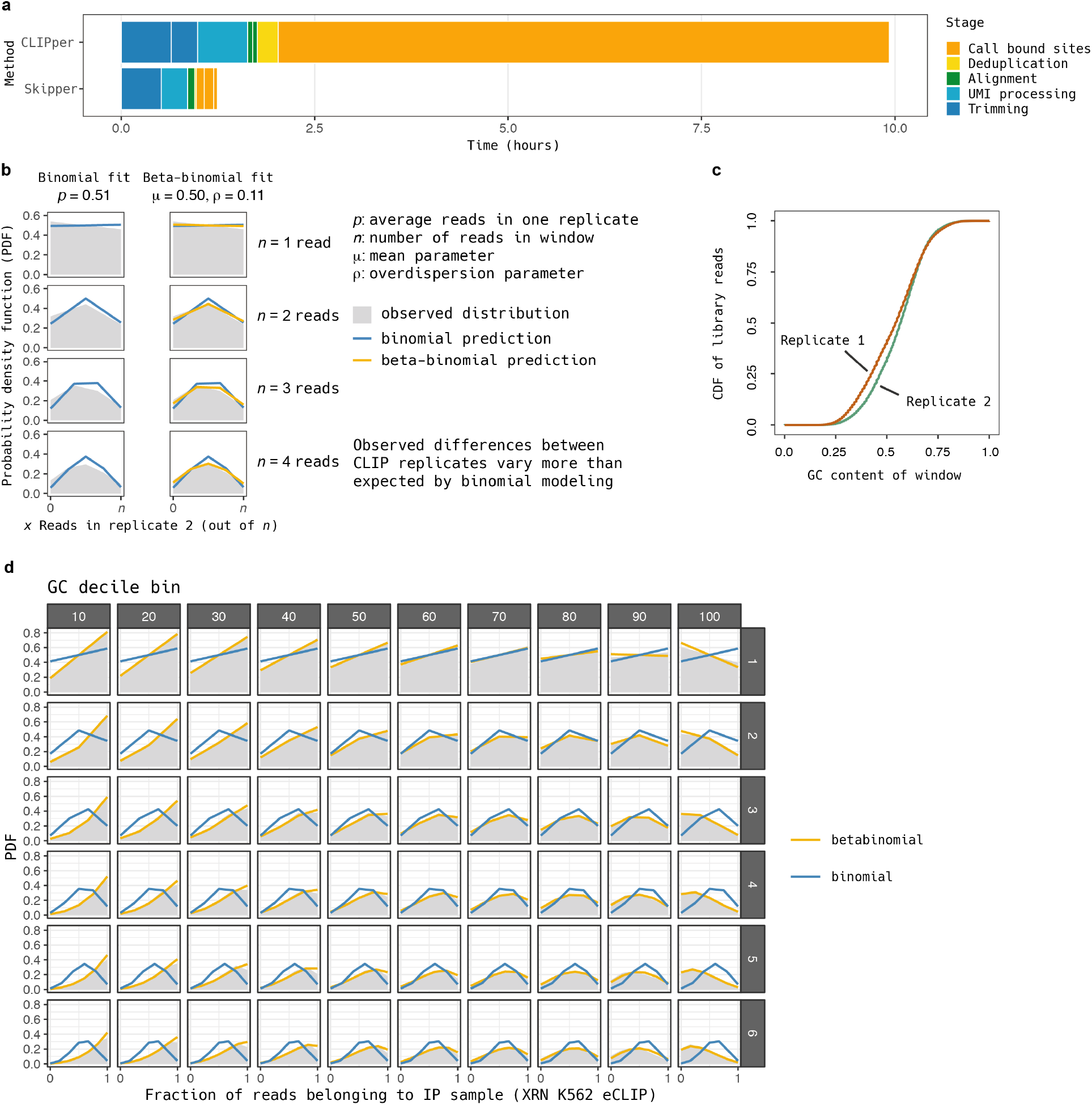
Quality control of Skipper model fitting. a) Average time spent processing Skipper and CLIPper with default options on a high performance computing system. b) Example fitting of binomial and beta-binomial models to two replicates of representative eCLIP data (PRPF8 eCLIP in HepG2 cells). c) Example GC bias across CLIP replicates for PRPF8 eCLIP in HepG2 cells. d) Example of beta-binomial modeling in the case of extreme GC content differences between CLIP and input (XRN eCLIP in K562).

**Supplementary Figure 2:**
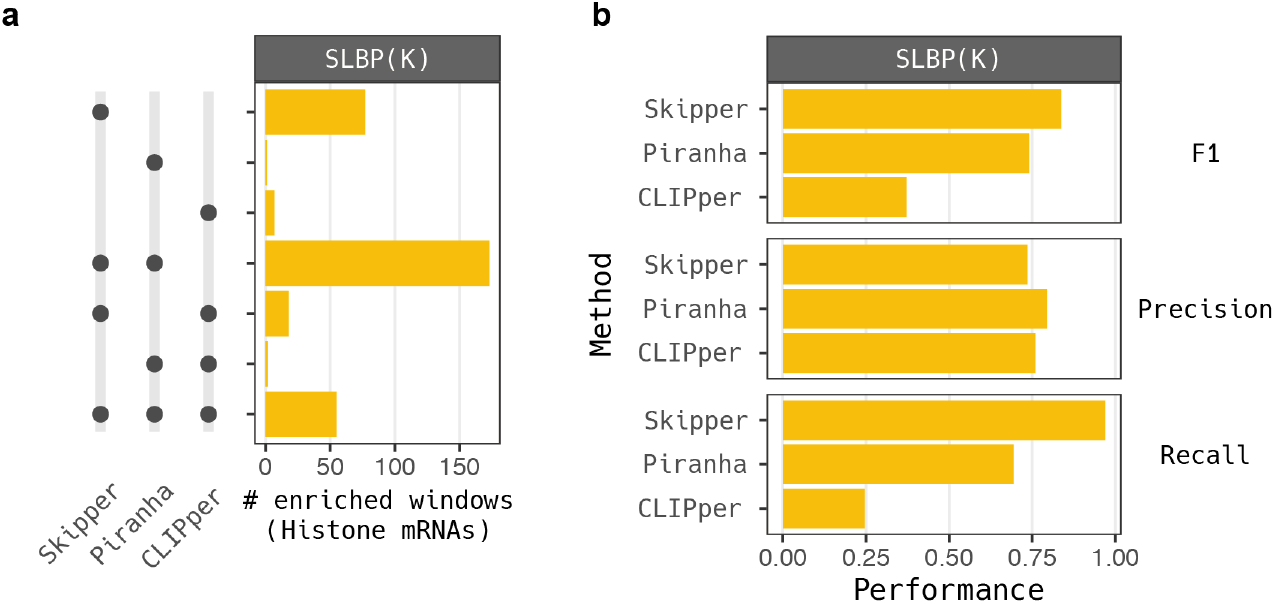
Binding site method performance on SLBP binding sites. a) Upset plot of the number of true positive enriched windows (histone mRNA targets) for Skipper, Piranha, and CLIPper. b) Precision, recall, and F1 measures for Skipper, Piranha, and CLIPper on SLBP.

**Supplementary Figure 3:**
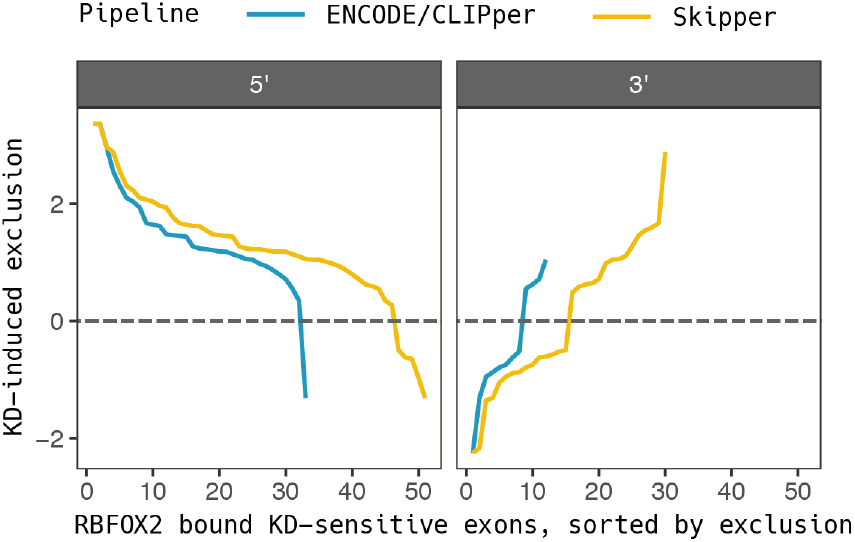
RBFOX2 binding near knockdown-sensitive alternative skipped exons. Knockdown-sensitive RBFOX2 exons sorted (x-axis) by average SepScore (y-axis) for proximity to both 5′ (left) and 3′ (right) splice sites.

**Supplementary Figure 4:**
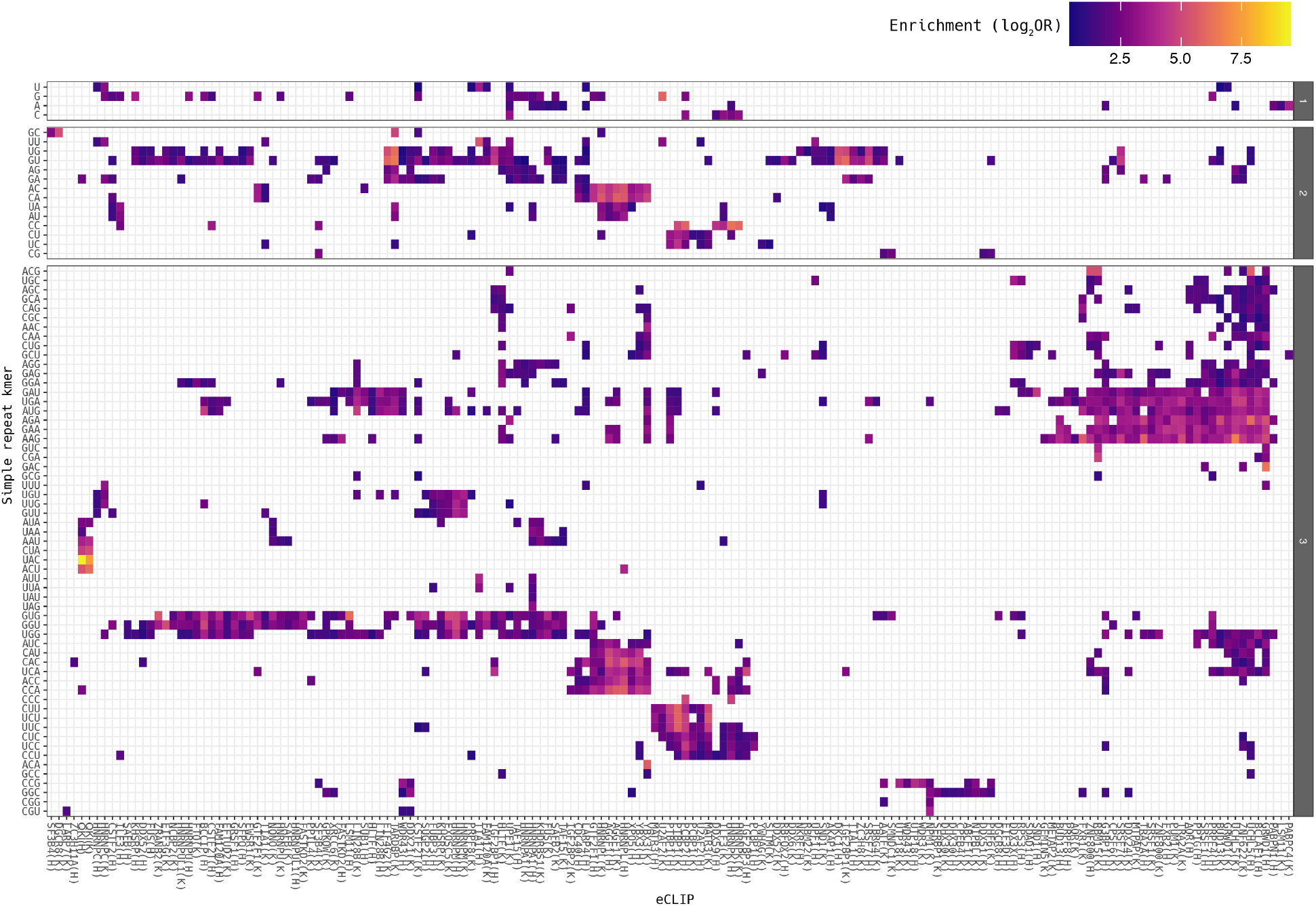
Simple repeat binding across eCLIPs. Colored tiles represent Skipper binding calls for mono-, di-, and trinucleotide simple repeats (y-axis) across CLIP experiments (x-axis). Heat denotes the level of enrichment.

**Supplementary Figure 5:**
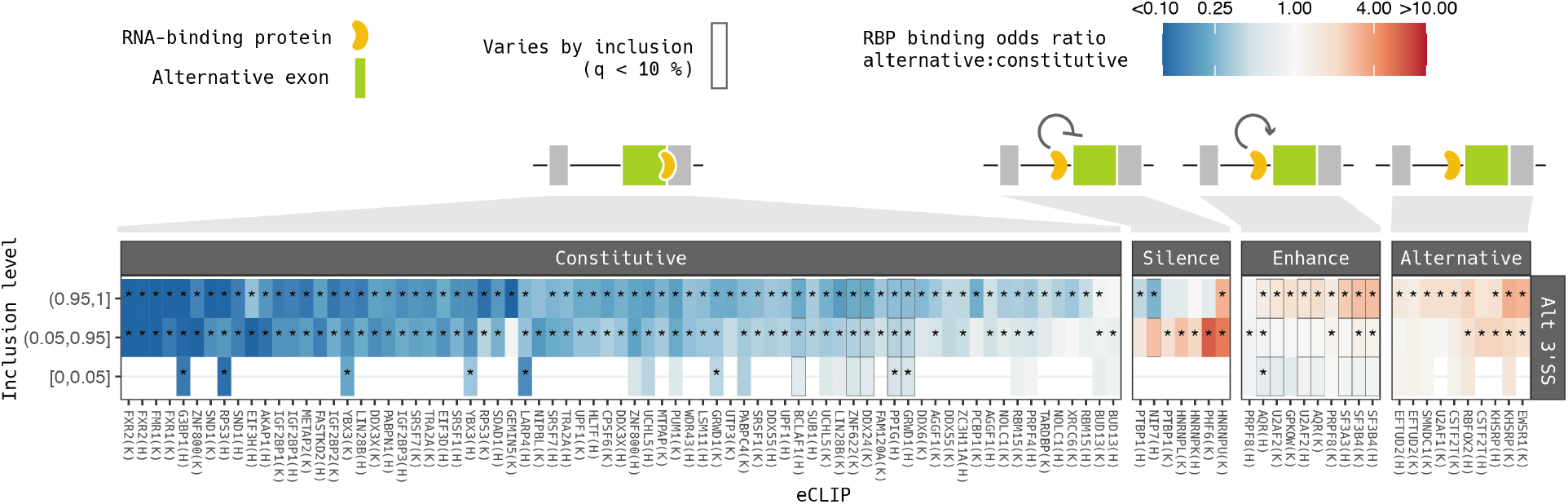
Archetypes of RNA-protein interactions near alternative 3′ splice sites. Patterns of binding and corresponding inclusion for RNA-binding proteins at alternative 3′ splice sites, as in Figure 4d.

**Supplementary Figure 6:**
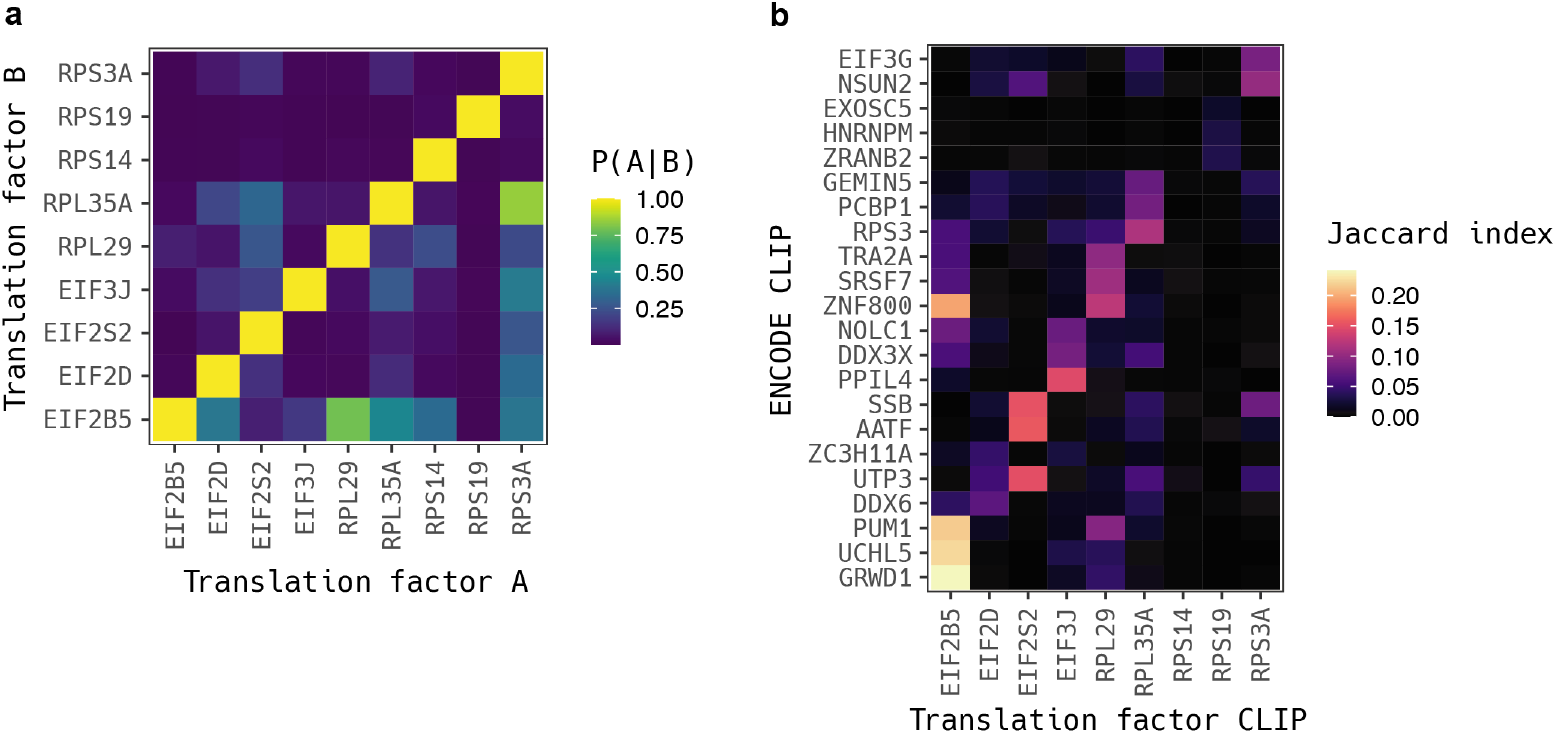
Overlapping enriched windows for translation factors. a) Fraction of enriched windows in the CLIP for the translation factor specified on the x-axis amongst enriched windows for the translation factor on the y-axis. b) Jaccard index of translation factor enriched windows intersected with previously published ENCODE CLIPs. The top three most similar CLIPs were selected for each translation factor.

**Supplementary Figure 7:**
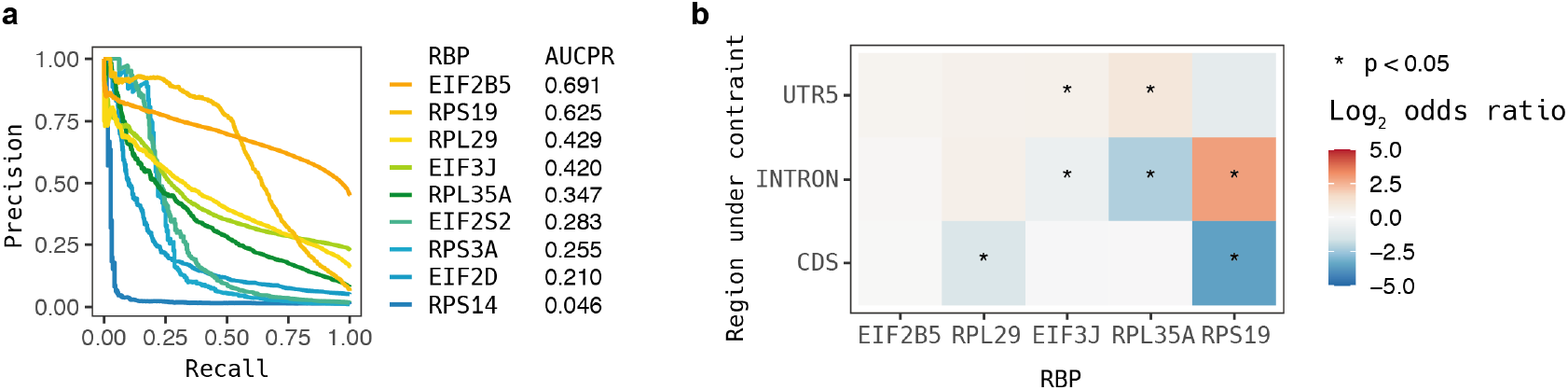
Properties of translation factors and binding sites under constraint. a) Cross-validation results on gapped kmer support vector machines trained on Skipper binding sites. High area under the precision recall curve (right) represents higher ability to predict translation factor occupancy using sequence alone, as seen for EIF2B5 and RPS19. b) Fisher’s Exact Test results for associating constrained binding with a disproportionate number of sites in the 5′ UTR, introns, or coding sequences. Regions with too few called binding sites were not considered.

**Supplementary Figure 8:**
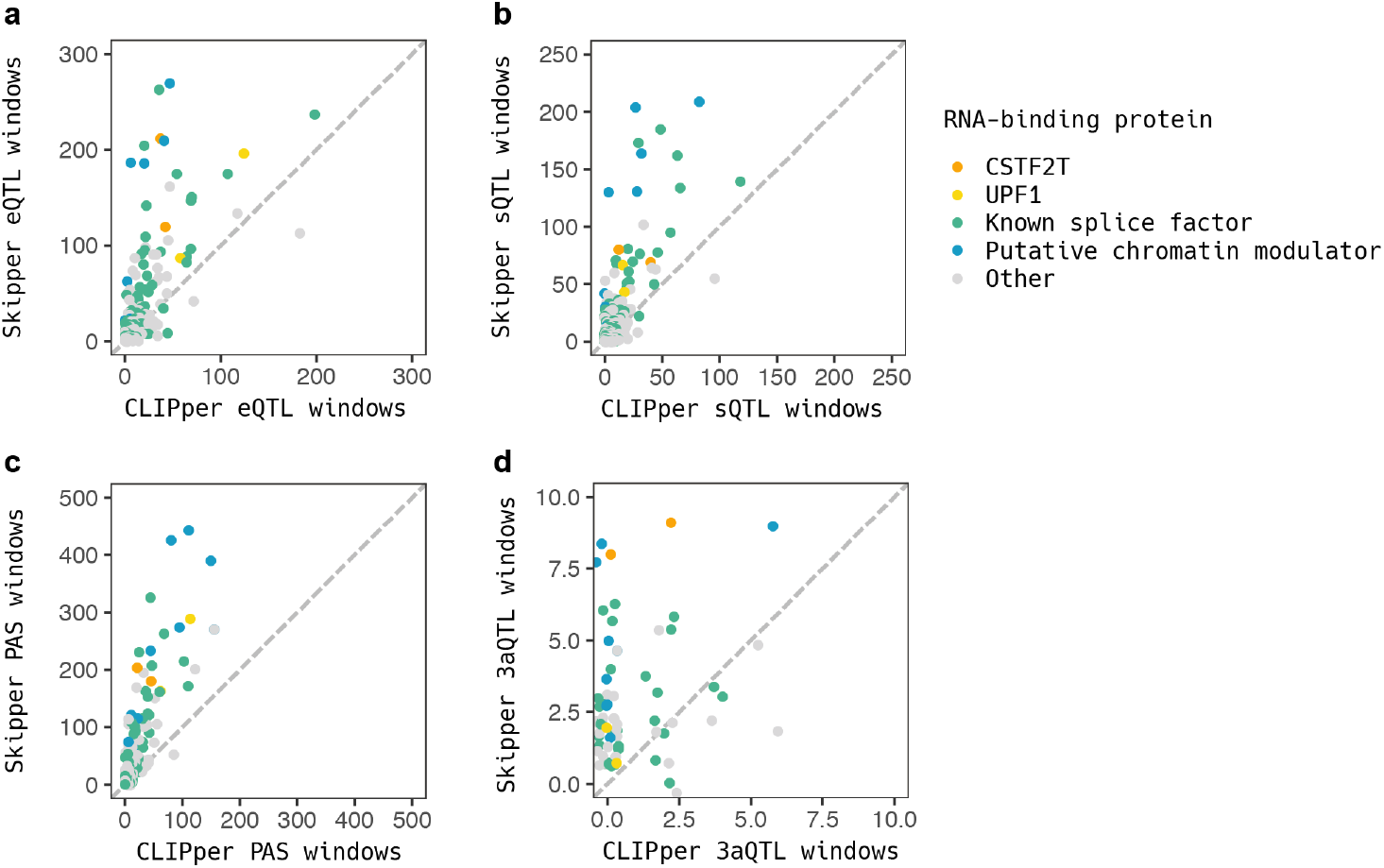
Intersection of Skipper binding sites and lead QTLs. Each point is one CLIP experiment comparing CLIPper overlaps (x-axis) to Skipper overlaps (y-axis). Putative chromatin modulators ZNF622, ZNF800, BCLAF1, YBX3, and GRWD1 are indicated in blue. Known splice factors are indicated in green. UPF1 and CSTF2T, two RBPs with abundant binding, are shown in yellow and orange. Overlaps are compared for a) lead eQTLs, b) lead sQTLs, c) polyadenylation sites, and d) lead 3′ alternate polyadenylation QTLs.

## References

1. Hentze, M. W., Castello, A., Schwarzl, T. & Preiss, T. A brave new world of RNA-binding proteins. Nat. Rev. Mol. Cell Biol. 19, 327–341 (2018).

2. Gerstberger, S., Hafner, M. & Tuschl, T. A census of human RNA-binding proteins. Nat. Rev. Genet. 15, 829–845 (2014).

3. Hafner, M. et al. CLIP and complementary methods. Nature Reviews Methods Primers 1, 1–23 (2021).

4. Wheeler, E. C., Van Nostrand, E. L. & Yeo, G. W. Advances and challenges in the detection of transcriptome-wide protein--RNA interactions. Wiley Interdiscip. Rev. RNA 9, e1436 (2018).

5. Van Nostrand, E. L. et al. Robust transcriptome-wide discovery of RNA-binding protein binding sites with enhanced CLIP (eCLIP). Nat. Methods 13, 508–514 (2016).

6. Uren, P. J. et al. Site identification in high-throughput RNA-protein interaction data. Bioinformatics 28, 3013–3020 (2012).

7. Katsantoni, M., van Nimwegen, E. & Zavolan, M. Improved analysis of (e)CLIP data with RCRUNCH yields a compendium of RNA-binding protein binding sites and motifs. bioRxiv (2022) doi:10.1101/2022.07.06.498949.

8. Feng, H. et al. Modeling RNA-Binding Protein Specificity In Vivo by Precisely Registering Protein-RNA Crosslink Sites. Mol. Cell 74, 1189–1204.e6 (2019).

9. Krakau, S., Richard, H. & Marsico, A. PureCLIP: capturing target-specific protein–RNA interaction footprints from single-nucleotide CLIP-seq data. Genome Biol. 18, 240 (2017).

10. Drewe-Boss, P., Wessels, H.-H. & Ohler, U. omniCLIP: probabilistic identification of protein-RNA interactions from CLIP-seq data. Genome Biol. 19, 183 (2018).

11. Zhang, Z. & Xing, Y. CLIP-seq analysis of multi-mapped reads discovers novel functional RNA regulatory sites in the human transcriptome. Nucleic Acids Res. 45, 9260–9271 (2017).

12. Van Nostrand, E. L. et al. Principles of RNA processing from analysis of enhanced CLIP maps for 150 RNA binding proteins. Genome Biol. 21, 90 (2020).

13. Uhl, M., Tran, V. D. & Backofen, R. Improving CLIP-seq data analysis by incorporating transcript information. BMC Genomics 21, 894 (2020).

14. Wagner, S. D. et al. Dose-Dependent Regulation of Alternative Splicing by MBNL Proteins Reveals Biomarkers for Myotonic Dystrophy. PLoS Genet. 12, e1006316 (2016).

15. Becker, W. R., Jarmoskaite, I., Vaidyanathan, P. P., Greenleaf, W. J. & Herschlag, D. Demonstration of protein cooperativity mediated by RNA structure using the human protein PUM2. RNA 25, 702–712 (2019).

16. Dassi, E. Handshakes and Fights: The Regulatory Interplay of RNA-Binding Proteins. Front Mol Biosci 4, 67 (2017).

17. Van Nostrand, E. L. et al. A large-scale binding and functional map of human RNA-binding proteins. Nature 583, 711–719 (2020).

18. Jarmoskaite, I. et al. A Quantitative and Predictive Model for RNA Binding by Human Pumilio Proteins. Mol. Cell 74, 966–981.e18 (2019).

19. Frankish, A. et al. GENCODE reference annotation for the human and mouse genomes. Nucleic Acids Res. 47, D766–D773 (2019).

20. Begg, B. E., Jens, M., Wang, P. Y., Minor, C. M. & Burge, C. B. Concentration-dependent splicing is enabled by Rbfox motifs of intermediate affinity. Nat. Struct. Mol. Biol. 27, 901–912 (2020).

21. Galarneau, A. & Richard, S. Target RNA motif and target mRNAs of the Quaking STAR protein. Nat. Struct. Mol. Biol. 12, 691–698 (2005).

22. Zhang, Y. et al. MATR3-antisense LINE1 RNA meshwork scaffolds higher-order chromatin organization. bioRxiv 2022.09.13.506124 (2022) doi:10.1101/2022.09.13.506124.

23. Xiong, F. et al. RNA m6A modification orchestrates a LINE-1–host interaction that facilitates retrotransposition and contributes to long gene vulnerability. Cell Res. 31, 861–885 (2021).

24. Attig, J. et al. Heteromeric RNP Assembly at LINEs Controls Lineage-Specific RNA Processing. Cell 174, 1067–1081.e17 (2018).

25. Liu, N. et al. Selective silencing of euchromatic L1s revealed by genome-wide screens for L1 regulators. Nature 553, 228–232 (2018).

26. Zarnack, K. et al. Direct competition between hnRNP C and U2AF65 protects the transcriptome from the exonization of Alu elements. Cell 152, 453–466 (2013).

27. Fasolo, F. et al. The RNA-binding protein ILF3 binds to transposable element sequences in SINEUP lncRNAs. FASEB J. 33, 13572–13589 (2019).

28. Thandapani, P., O’Connor, T. R., Bailey, T. L. & Richard, S. Defining the RGG/RG motif. Mol. Cell 50, 613–623 (2013).

29. Yagi, R., Miyazaki, T. & Oyoshi, T. G-quadruplex binding ability of TLS/FUS depends on the β-spiral structure of the RGG domain. Nucleic Acids Res. 46, 5894–5901 (2018).

30. Masuzawa, T. & Oyoshi, T. Roles of the RGG Domain and RNA Recognition Motif of Nucleolin in G-Quadruplex Stabilization. ACS Omega 5, 5202–5208 (2020).

31. Lee, D. S. M., Ghanem, L. R. & Barash, Y. Integrative analysis reveals RNA G-quadruplexes in UTRs are selectively constrained and enriched for functional associations. Nat. Commun. 11, 527 (2020).

32. Ruggiero, E. et al. Fused in Liposarcoma Protein, a New Player in the Regulation of HIV-1 Transcription, Binds to Known and Newly Identified LTR G-Quadruplexes. ACS Infect Dis 8, 958–968 (2022).

33. Butovskaya, E., Heddi, B., Bakalar, B., Richter, S. N. & Phan, A. T. Major G-quadruplex form of HIV-1 LTR reveals a (3 + 1) folding topology containing a stem-loop. J. Am. Chem. Soc. 140, 13654–13662 (2018).

34. Jaganathan, K. et al. Predicting Splicing from Primary Sequence with Deep Learning. Cell 176, 535–548.e24 (2019).

35. Garrido-Martín, D., Borsari, B., Calvo, M., Reverter, F. & Guigó, R. Identification and analysis of splicing quantitative trait loci across multiple tissues in the human genome. Nat. Commun. 12, 727 (2021).

36. Qi, T. et al. Genetic control of RNA splicing and its distinct role in complex trait variation. Nat. Genet. (2022) doi:: 10.1038/s41588-022-01154-4.

37. Li, Y. I. et al. Annotation-free quantification of RNA splicing using LeafCutter. Nat. Genet. 50, 151–158 (2018).

38. Yang, E.-W. et al. Allele-specific binding of RNA-binding proteins reveals functional genetic variants in the RNA. Nat. Commun. 10, 1338 (2019).

39. Ashburner, M. et al. Gene Ontology: tool for the unification of biology. Nat. Genet. 25, 25–29 (2000).

40. Subramanian, A. et al. Gene set enrichment analysis: a knowledge-based approach for interpreting genome-wide expression profiles. Proc. Natl. Acad. Sci. U. S. A. 102, 15545–15550 (2005).

41. Liberzon, A. et al. The Molecular Signatures Database (MSigDB) hallmark gene set collection. Cell Syst 1, 417–425 (2015).

42. Adamson, S. I., Zhan, L. & Graveley, B. R. Functional characterization of splicing regulatory elements. bioRxiv (2021) doi:: 10.1101/2021.05.14.444228.

43. Rambout, X., Dequiedt, F. & Maquat, L. E. Beyond Transcription: Roles of Transcription Factors in Pre-mRNA Splicing. Chem. Rev. 118, 4339–4364 (2018).

44. Ma, J. et al. The requirement of the DEAD-box protein DDX24 for the packaging of human immunodeficiency virus type 1 RNA. Virology 375, 253–264 (2008).

45. Zeng, M., Zhu, L., Li, L. & Kang, C. miR-378 suppresses the proliferation, migration and invasion of colon cancer cells by inhibiting SDAD1. Cell. Mol. Biol. Lett. 22, 12 (2017).

46. Thul, P. J. et al. A subcellular map of the human proteome. Science 356, (2017).

47. Samarsky, D. A., Fournier, M. J., Singer, R. H. & Bertrand, E. The snoRNA box C/D motif directs nucleolar targeting and also couples snoRNA synthesis and localization. EMBO J. 17, 3747–3757 (1998).

48. Young, D. J., Meydan, S. & Guydosh, N. R. 40S ribosome profiling reveals distinct roles for Tma20/Tma22 (MCT-1/DENR) and Tma64 (eIF2D) in 40S subunit recycling. Nat. Commun. 12, 2976 (2021).

49. Lek, M. et al. Analysis of protein-coding genetic variation in 60,706 humans. Nature 536, 285–291 (2016).

50. Park, C. Y. et al. Genome-wide landscape of RNA-binding protein target site dysregulation reveals a major impact on psychiatric disorder risk. Nat. Genet. 53, 166–173 (2021).

51. Lee, D. et al. A method to predict the impact of regulatory variants from DNA sequence. Nat. Genet. 47, 955–961 (2015).

52. Ghanbari, M. & Ohler, U. Deep neural networks for interpreting RNA-binding protein target preferences. Genome Res. 30, 214–226 (2020).

53. Denichenko, P. et al. Specific inhibition of splicing factor activity by decoy RNA oligonucleotides. Nat. Commun. 10, 1590 (2019).

54. Arandel, L. et al. Reversal of RNA toxicity in myotonic dystrophy via a decoy RNA-binding protein with high affinity for expanded CUG repeats. Nat Biomed Eng 6, 207–220 (2022).

55. Jackson, A. L. et al. Widespread siRNA “off-target” transcript silencing mediated by seed region sequence complementarity. RNA 12, 1179–1187 (2006).

56. Zhang, X. et al. Mechanisms and Functions of Long Non-Coding RNAs at Multiple Regulatory Levels. Int. J. Mol. Sci. 20, (2019).

57. Rom, A. et al. Regulation of CHD2 expression by the Chaserr long noncoding RNA gene is essential for viability. Nat. Commun. 10, 5092 (2019).

58. Da Costa, L., Leblanc, T. & Mohandas, N. Diamond-Blackfan anemia. Blood 136, 1262–1273 (2020).

59. Moras, M., Lefevre, S. D. & Ostuni, M. A. From Erythroblasts to Mature Red Blood Cells: Organelle Clearance in Mammals. Front. Physiol. 8, 1076 (2017).

60. Mortensen, M. et al. Loss of autophagy in erythroid cells leads to defective removal of mitochondria and severe anemia in vivo. Proc. Natl. Acad. Sci. U. S. A. 107, 832–837 (2010).

61. Doulatov, S. et al. Drug discovery for Diamond-Blackfan anemia using reprogrammed hematopoietic progenitors. Sci. Transl. Med. 9, (2017).

62. Patro, R., Duggal, G., Love, M. I., Irizarry, R. A. & Kingsford, C. Salmon provides fast and bias-aware quantification of transcript expression. Nat. Methods 14, 417–419 (2017).

63. Jiang, H., Lei, R., Ding, S.-W. & Zhu, S. Skewer: a fast and accurate adapter trimmer for next-generation sequencing paired-end reads. BMC Bioinformatics 15, 182 (2014).

64. Chen, S., Zhou, Y., Chen, Y. & Gu, J. fastp: an ultra-fast all-in-one FASTQ preprocessor. Bioinformatics 34, i884–i890 (2018).

65. Dobin, A. et al. STAR: ultrafast universal RNA-seq aligner. Bioinformatics 29, 15–21 (2013).

66. Liu, D. Algorithms for efficiently collapsing reads with Unique Molecular Identifiers. PeerJ 7, e8275 (2019).

67. Li, H. et al. The Sequence Alignment/Map format and SAMtools. Bioinformatics 25, 2078–2079 (2009).

68. Quinlan, A. R. & Hall, I. M. BEDTools: a flexible suite of utilities for comparing genomic features. Bioinformatics 26, 841–842 (2010).

69. Yee, T. W. Vector Generalized Linear and Additive Models. (Springer New York, 2015).

70. Krijthe. Rtsne: T-distributed stochastic neighbor embedding using Barnes-Hut implementation. R package version 0.13, URL https://github.com/jkrijthe (2015).

71. Rasheedi, Shun, Serrao & Sowd. The cleavage and polyadenylation specificity factor 6 (CPSF6) subunit of the capsid-recruited pre-messenger RNA cleavage factor I (CFIm) complex mediates …. Boll. Soc. Ital. Biol. Sper.

72. Aznarez, Barash, Shai, He & Zielenski. A systematic analysis of intronic sequences downstream of 5′ splice sites reveals a widespread role for U-rich motifs and TIA1/TIAL1 proteins in alternative splicing …. Genome.

73. Blue, S. M. et al. Transcriptome-wide identification of RNA-binding protein binding sites using seCLIP-seq. Nat. Protoc. 17, 1223–1265 (2022).

74. Anger, A. M. et al. Structures of the human and Drosophila 80S ribosome. Nature 497, 80–85 (2013).

75. Lee, D. LS-GKM: a new gkm-SVM for large-scale datasets. Bioinformatics 32, 2196–2198 (2016).

76. Sing, T., Sander, O., Beerenwinkel, N. & Lengauer, T. ROCR: visualizing classifier performance in R. Bioinformatics 21, 3940–3941 (2005).

77. Li, L. et al. An atlas of alternative polyadenylation quantitative trait loci contributing to complex trait and disease heritability. Nat. Genet. 53, 994–1005 (2021).

78. Mittleman, B. E. et al. Alternative polyadenylation mediates genetic regulation of gene expression. Elife 9, (2020).

